# Coordinated inheritance of extrachromosomal DNA species in human cancer cells

**DOI:** 10.1101/2023.07.18.549597

**Authors:** King L. Hung, Matthew G. Jones, Ivy Tsz-Lo Wong, Joshua T. Lange, Jens Luebeck, Elisa Scanu, Britney Jiayu He, Lotte Brückner, Rui Li, Rocío Chamorro González, Rachel Schmargon, Jan R. Dörr, Julia A. Belk, Vineet Bafna, Benjamin Werner, Weini Huang, Anton G. Henssen, Paul S. Mischel, Howard Y. Chang

## Abstract

The chromosomal theory of inheritance has dominated human genetics, including cancer genetics. Genes on the same chromosome segregate together while genes on different chromosomes assort independently, providing a fundamental tenet of Mendelian inheritance. Extrachromosomal DNA (ecDNA) is a frequent event in cancer that drives oncogene amplification, dysregulated gene expression and intratumoral heterogeneity, including through random segregation during cell division. Distinct ecDNA sequences, herein termed ecDNA species, can co-exist to facilitate intermolecular cooperation in cancer cells. However, how multiple ecDNA species within a tumor cell are assorted and maintained across somatic cell generations to drive cancer cell evolution is not known. Here we show that cooperative ecDNA species can be coordinately inherited through mitotic co-segregation. Imaging and single-cell analyses show that multiple ecDNAs encoding distinct oncogenes co-occur and are correlated in copy number in human cancer cells. EcDNA species are coordinately segregated asymmetrically during mitosis, resulting in daughter cells with simultaneous copy number gains in multiple ecDNA species prior to any selection. Computational modeling reveals the quantitative principles of ecDNA co-segregation and co-selection, predicting their observed distributions in cancer cells. Finally, we show that coordinated inheritance of ecDNAs enables co-amplification of specialized ecDNAs containing only enhancer elements and guides therapeutic strategies to jointly deplete cooperating ecDNA oncogenes. Coordinated inheritance of ecDNAs confers stability to oncogene cooperation and novel gene regulatory circuits, allowing winning combinations of epigenetic states to be transmitted across cell generations.

## INTRODUCTION

Oncogene amplification drives cancer development by increasing the copies of genetic sequences that encode oncogene products. Oncogenes are frequently amplified on megabase-sized circular extrachromosomal DNA (ecDNA), which is detected in half of human cancer types^1^. Patients with tumors containing ecDNA have shorter survival than those with tumors harboring other types of focal amplification^2, 3^, suggesting that ecDNA-driven oncogene amplification may make tumors more aggressive. This aggressive behavior may be attributed to the ability of ecDNA-containing cancer cells to rapidly adapt to selective pressures. EcDNA is replicated in each cell cycle and transmitted through cell division. However, as it lacks centromeres, ecDNA segregates randomly to daughter cells during mitosis, leading to copy number heterogeneity^4–6^. This copy number heterogeneity enables more rapid changes to the DNA contents of cells and supports adaptation to new selective pressures such as metabolic stress and drug treatment^4, 7, 8^.

EcDNAs exhibit a remarkable level of genetic sequence diversity in parallel to their copy number diversity^3, 9–13^. First, multiple ecDNAs that are originally derived from different chromosomal loci can co-exist in the same cancer cells. These ecDNAs can congregate in micron-sized hubs in the nucleus and enable intermolecular gene activation, where enhancer elements on one ecDNA molecule can activate coding sequences on another ecDNA^9^. Second, ecDNAs harbor clustered somatic mutations that suggest APOBEC3-mediated mutagenesis^14, 15^. These unique, subclonal mutations within oncogenes or other functional elements on ecDNAs increase the diversity of ecDNA sequence and function with potential impacts on tumor evolution^8, 10, 15^. Third, ecDNAs can contain complex structural rearrangements, resulting from recombination of genomic sequences originating from various genomic sites or different chromosomes^2, 9–13, 16^. These complex circularization events can give rise to diverse ecDNA species co-existing in a cancer cell population, including ecDNAs with distinct oncogene loci^3, 9–13^ or ecDNAs encompassing only enhancers or oncogene coding sequences^10^.

EcDNAs may represent specialized molecules that cooperate to increase cancer cell fitness. It has been reported that additional ecDNA species can form after recurrence or drug treatment of ecDNA-carrying cancers^10–12;^ the original ecDNA amplicons were retained in the recurrent cancer cells in these studies, suggesting that multiple ecDNA species may arise independently and provide fitness advantages to cancer cells. Heterogeneous ecDNA species co-occurring in the same cell can contain distinct oncogenes^9–13^. These ecDNAs carrying oncogenes as well as non-coding regulatory elements can interact with each other and with chromosomes in an intermolecular, combinatorial manner to promote gene expression^9, 17, 18^. These observations suggest that the co-occurrence of multiple ecDNA sequences in a cell may have combinatorial and synergistic effects on transcriptional programs.

This diversity of ecDNA genetic sequences in a cancer cell population raises the following questions: 1) How are heterogeneous ecDNA species distributed in a cell population? 2) As ecDNAs are segregated unequally during mitosis, how are these mixtures of ecDNAs inherited by daughter cells? 3) How do the dynamics of multiple ecDNA species affect cancer evolution? To address these questions, we used a combination of image analysis, single-cell and bulk sequencing, and computational modeling to elucidate the principles and consequences of ecDNA co-evolution in cancer.

## RESULTS

### EcDNAs encoding distinct oncogenes co-occur in human cancers and single cancer cells

To interrogate how frequently ecDNA molecules with distinct sequences co-exist in the same tumors, we first analyzed the structures of focal amplifications in whole-genome sequencing (WGS) data from 1513 patient tumors from The Cancer Genome Atlas (TCGA)^2^. 289 of 1513 patient tumors contained ecDNA, carrying coding sequences of well-characterized oncogenes such as *EGFR*, *MDM2* and *CDK4*^1, 2^ (**Figure 1a,b**; Methods). Of tumors that contained ecDNA, more than 25% (81 samples) contained two or more ecDNA species in the same tumor (**Figure 1a**). Many of these ecDNA species were amplified at high copy numbers and contained canonical oncogenes (**Figure 1b**). This result supports the idea that heterogeneous ecDNA sequences can be found in the same tumor and their co-occurrence may provide distinct selective advantages. As we only considered highly abundant and genomically non-overlapping ecDNA sequences as distinct species, this analysis likely underestimates the true diversity of ecDNA species.

**Figure 1.**
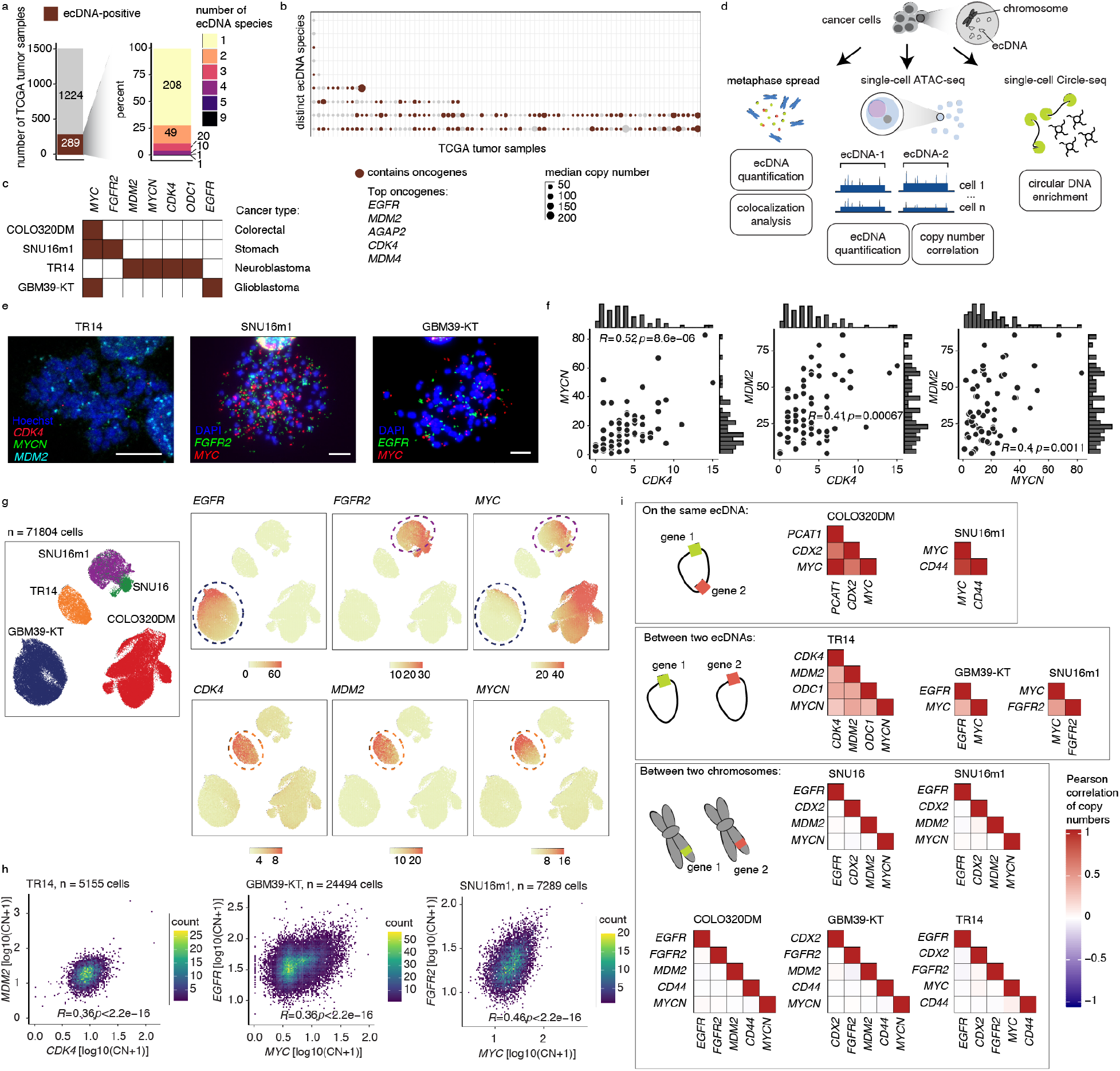
EcDNA species encoding distinct oncogene sequences are correlated in individual cancer cells. **(a)** Summary of ecDNA-positive tumors (left) and number of ecDNA species (right) identified in TCGA tumor samples. **(b)** Median copy numbers and oncogene statuses of distinct ecDNA species in TCGA tumors identified to have more than one ecDNA species. **(c)** A panel of cell lines with known oncogene sequences on ecDNA. **(d)** Schematic of ecDNA analyses using three orthogonal approaches: metaphase spread, scATAC-seq, and scCircle-seq. **(e)** Representative DNA FISH images of metaphase spreads with FISH probes targeting various oncogene sequences as indicated. Scale bars are 10 µm. **(f)** Oncogene copy number scatter plots, histograms and Spearman correlations between pairs of oncogenes in TR14 cells. **(g)** UMAP from scATAC-seq showing cell line annotations (left) and copy number calculations of indicated oncogenes (right panels). **(h)** Density scatter plots of log-transformed oncogene copy numbers and Pearson correlations between pairs of oncogenes in the indicated cell lines. **(i)** Pearson correlation heatmaps of gene pairs on the same ecDNA (top), between two ecDNAs (middle), and between two chromosomes (bottom).

The frequent co-amplification of distinct ecDNA species in tumors raised the question of whether multiple ecDNA species can co-occur in the same cells. To address this question, we examined a panel of cancer cell line and neurosphere models representing colorectal cancer, neuroblastoma, glioblastoma, and stomach cancer that have previously been characterized as containing ecDNA species with distinct amplified oncogenes (**Figure 1c**). Using DNA fluorescent *in situ* hybridization (FISH) of metaphase-spread chromosomes, we validated three cell lines previously characterized to contain multiple amplified oncogenes on ecDNA: the monoclonal SNU16m1 stomach cancer line contained *FGFR2* and *MYC* ecDNAs, the TR14 neuroblastoma cell line contained *MYCN*, *CDK4*, *ODC1* and *MDM2* ecDNAs, and the GBM39-KT glioblastoma neurospheres, a subline of the well-characterized GBM39 culture with *EGFR* amplified on ecDNA that also developed *MYC* amplification on ecDNA (**Figure 1d,e**). Importantly, metaphase FISH confirmed that the vast majority of individual cells have very little overlap in FISH signals from distinct oncogenes, showing that they are not covalently linked on the same ecDNA molecule and therefore are expected to be inherited independently from one another in dividing cancer cells (**Figure 1e, Extended Data Figure 1a-c**).

We next examined distributions of ecDNA copy numbers in single cells using three orthogonal methods (**Figure 1d**): 1) metaphase chromosome spreading followed by DNA FISH; 2) isolation of single nuclei followed by droplet-based single-cell assay for transposase-accessible chromatin using sequencing (scATAC-seq) and RNA sequencing; and 3) enrichment and sequencing of ecDNAs in individual cells via exonuclease digestion and rolling circle amplification^19, 20^ (single-cell Circle-seq; scCircle-seq; Methods). Remarkably, in the cell lines with distinct ecDNA species, FISH imaging revealed that pairs of ecDNA species had significantly correlated copy numbers (Spearman correlation *R* = 0.39-0.52, *p* < 0.01 in all cases; **Figure 1f, Extended Data Figure 1c**). Next, we assessed the significance of these correlations in a larger cell population by adapting a copy number quantification method for genomic background coverage from scATAC-seq data^9, 21, 22^ to calculate ecDNA copy numbers in these cell lines in an integrated analysis of 71,804 cells (**Figure 1d,g, Extended Data Figure 2a**; Methods). Strikingly, we observed positive correlations between distinct ecDNA species in each of the three cell lines with multiple ecDNA species (**Figure 1h,I, Extended Data Figure 2b**; Pearson correlation *R* = 0.26-0.46, *p* < 1e10^-15^ in all cases). As expected, genic sequences that are covalently linked on the same ecDNA molecule (as demonstrated by isolation from the same molecular size fractions by CRISPR-CATCH^10^; **Extended Data Figure 2c**) showed strong copy number correlation in this analysis, validating this approach for measuring distributions of ecDNA molecules in a cell population (**Figure 1i, Extended Data Figure 2b**). EcDNA copy numbers were positively correlated with the RNA expression of the correspondingly amplified oncogenes, validating that the copies of ecDNA species drive transcriptional outcomes (**Extended Data Figure 2d**). Importantly, we did not observe copy number correlations between gene pairs located on different chromosomes, suggesting that this relationship between different ecDNA species cannot simply be explained by differences in sequencing depth or sequencing quality between single cells (**Figure 1i, Extended Data Figure 2b**). Finally, single cell Circle-seq confirmed co-enrichment of the *MYCN*, *MDM2* and *CDK4* ecDNA species in individual TR14 neuroblastoma cells (**Extended Data Figure 2e**). These results show that distinct ecDNA species tend to co-occur with correlated copy numbers far more than expected by chance in human cancer cells.

**Figure 2.**
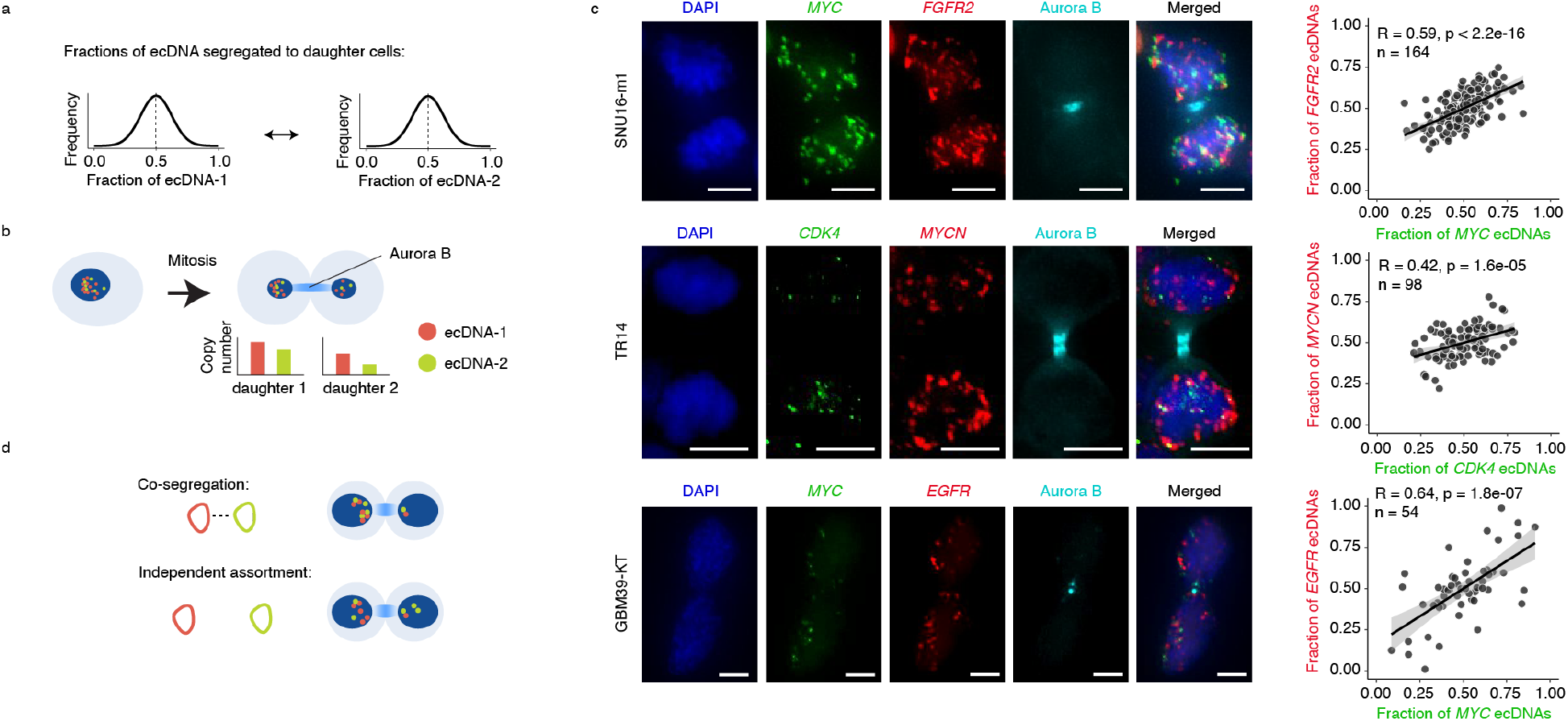
Distinct ecDNA species are co-segregated into daughter cells during mitosis. **(a)** Copy numbers of individual ecDNA species in a daughter cell after segregation are predicted to follow a Gaussian distribution due to random segregation. **(b)** Pairs of daughter cells undergoing mitosis are identified by immunofluorescence for Aurora kinase B (Aurora B). Individual ecDNA species are visualized using FISH probes targeting specific oncogene sequences for copy number quantification. **(c)** Representative images of pairs of daughter cells undergoing mitosis (left). Scatter plots showing Pearson correlations of per-cell ecDNA contents containing the indicated oncogene sequences in daughter cells (right). Scale bars are 5 µm. **(d)** Schematic of co-segregation and independent assortment of multiple ecDNA species in mitosis.

### Distinct ecDNA species co-segregate to daughter cells during mitosis

In principle, our observations of co-occurrence and correlation of two distinct ecDNA species can be the result of 1) co-selection of both species, given that both species provide fitness advantages and/or engage in synergistic intermolecular interactions, or 2) co-segregation of both species into daughter cells during cell division. As different ecDNA species can carry different oncogenes and mixed ecDNAs can interact with each other to increase gene expression^9, 18^, co-selection can reasonably explain co-occurrence of ecDNA species. However, unlike the faithful segregation of chromosomes, ecDNAs lack centromeres and are randomly inherited during mitosis^4–6^. Therefore, it is unclear how a population of ecDNA species and their cooperative interactions are preserved over successive cell divisions (**Figure 2a**).

To address this question, we assessed the distribution of multiple ecDNA species during each single cell division. Specifically, we used DNA FISH combined with immunofluorescence staining for Aurora kinase B, a component of the mitotic midbody, to quantify copy numbers of ecDNA inherited among pairs of daughter cells undergoing mitosis^4, 23^ (**Figure 2b**). Interestingly, in all three cancer cell types that showed copy number correlations of multiple ecDNAs at the population level, we observed significant co-segregation of distinct ecDNA species to daughter cells in mitosis (*R* = 0.42-0.64, *p* < 1e10^-4^ in each case; **Figure 2c**; Methods), contrary to the Mendelian principle of independent assortment. In other words, the daughter cell that inherits more copies of ecDNA species 1 tends to inherit more copies of species 2, and vice versa. Close inspection of the FISH images suggested that the co-segregating ecDNAs remain largely distinct despite being in the same nuclei. Computer simulations of segregating ecDNAs showed that this correlation of ecDNA species in daughter cells is far greater than expected from random segregation and scales linearly with the level of co-segregation of ecDNAs (**Extended Data Figure 3**; Methods). Together, these data show that while individual ecDNAs segregate in a random manner^4, 6^, collectives of ecDNA species may co-segregate during mitosis (**Figure 2d**).

**Figure 3.**
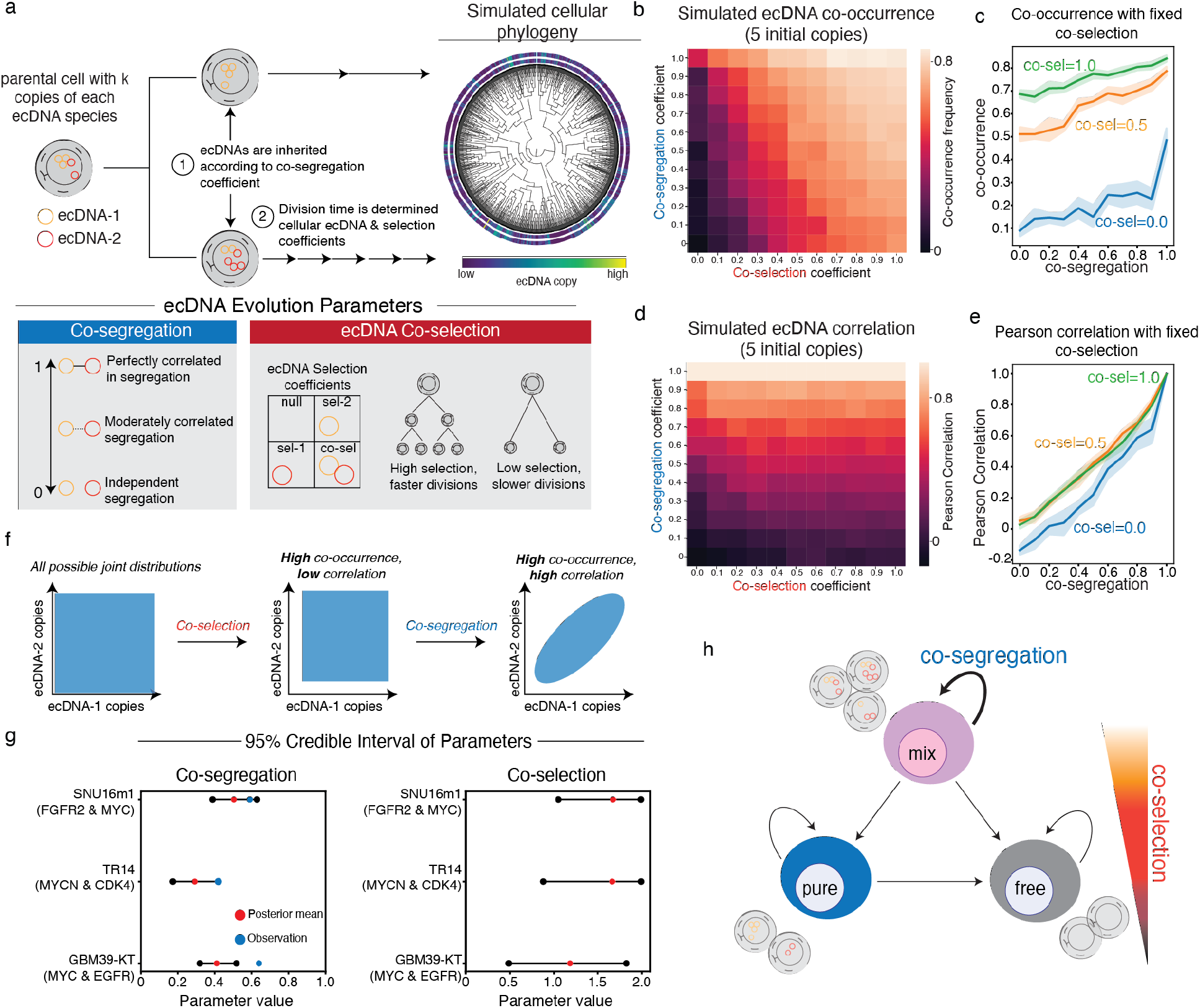
Evolutionary modeling of ecDNA dynamics reveals the principles of ecDNA co-inheritance. **(a)** The evolutionary modeling framework used in this study. Cancer populations are simulated starting from a single parent cell carrying *k* different ecDNA species (here, *k=2*) with user-defined initial copy-numbers. Cells divide according to a fitness function, parameterized by user-defined selection coefficients. During cell division, ecDNA is inherited according to a co-segregation coefficient. **(b-e)** Summary statistics of one-million-cell populations and 10 replicates across varying co-selection and co-segregation coefficients beginning with a parental cell with 5 copies of each ecDNA species. The average frequency of cells carrying both ecDNA species **(b)** and the Pearson correlation of ecDNA copy number within cells **(d)** are shown across all simulations. The mean frequency of cells carrying both ecDNA species **(c)** and Pearson’s correlation of ecDNA copy number within cells **(e)** are shown as a function of co-segregation level for fixed levels of co-selection: 0.0, 0.5, and 1.0 (shaded area represents the 95% confidence interval across the experimental replicates). Selection acting on cells carrying one but not both ecDNAs is maintained at 0.2 and selection acting on cells without either ecDNA is maintained at 0.0 across all simulations. **(f)** Schematic representation of the effects of co-selection and co-segregation on the joint distribution of ecDNA copy numbers in cancer cells. **(g)** 95% credible interval for inferred co-segregation and co-selection values for SNU16m1, TR14, and GBM39-KT cell lines. **(h)** Conceptual summary of ecDNA co-evolutionary dynamics.

### Evolutionary modeling infers the principles of ecDNA co-assortment

Next, we used evolutionary modeling to assess the contributions of co-selection and co-segregation in shaping the patterns of ecDNA co-assortment. Similar to previous work^4^, we implemented an individual-based, forward-time evolutionary algorithm to study ecDNA evolution in a growing tumor population (**Figure 3a**; Methods). This model is instantiated with a single parent cell carrying two distinct ecDNA species with the same copy number. The simulation proceeds by choosing cells to divide (or, optionally, die) according to a “fitness” function that determines the birth rate of a cell based on the presence of each ecDNA species. During cell division, ecDNA copies are inherited amongst daughter cells according to a “co-segregation” parameter: a value of 0 indicates independent, random segregation and a value of 1 indicates perfectly correlated segregation.

Under fixed selection for individual ecDNA species, we first studied how varying co-segregation and co-selection parameters affected the co-assortment of two ecDNA species in simulations of one million tumor cells (**Figure 3b-e**; **Extended Data Figure 4**). Though ecDNA copy numbers rose in all simulations (**Extended Data Figure 4a**), these simulations revealed specific principles of ecDNA co-evolution arising from co-segregation and co-selection: 1) co-occurrence (the frequency of cells carrying both ecDNA species) was predominantly driven by co-selection pressure that acts over multiple generations to select for cells carrying both ecDNA species (**Figure 3b**), though both co-segregation and co-selection could synergize to achieve high-levels of co-occurrence (**Figure 3c**); and 2) copy-number correlation in cells appeared to be entirely driven by co-segregation alone, where proportional amounts of ecDNA copies can be inherited in a single cell division (**Figure 3d,e**). These trends were validated using an alternative model of ecDNA evolution (**Extended Data Figure 5**). Our simulations additionally suggested that ecDNA co-occurrence may be longer-lasting once the cancer cell population reaches high copy numbers (**Extended Data Figure 4b-d**).

**Figure 4.**
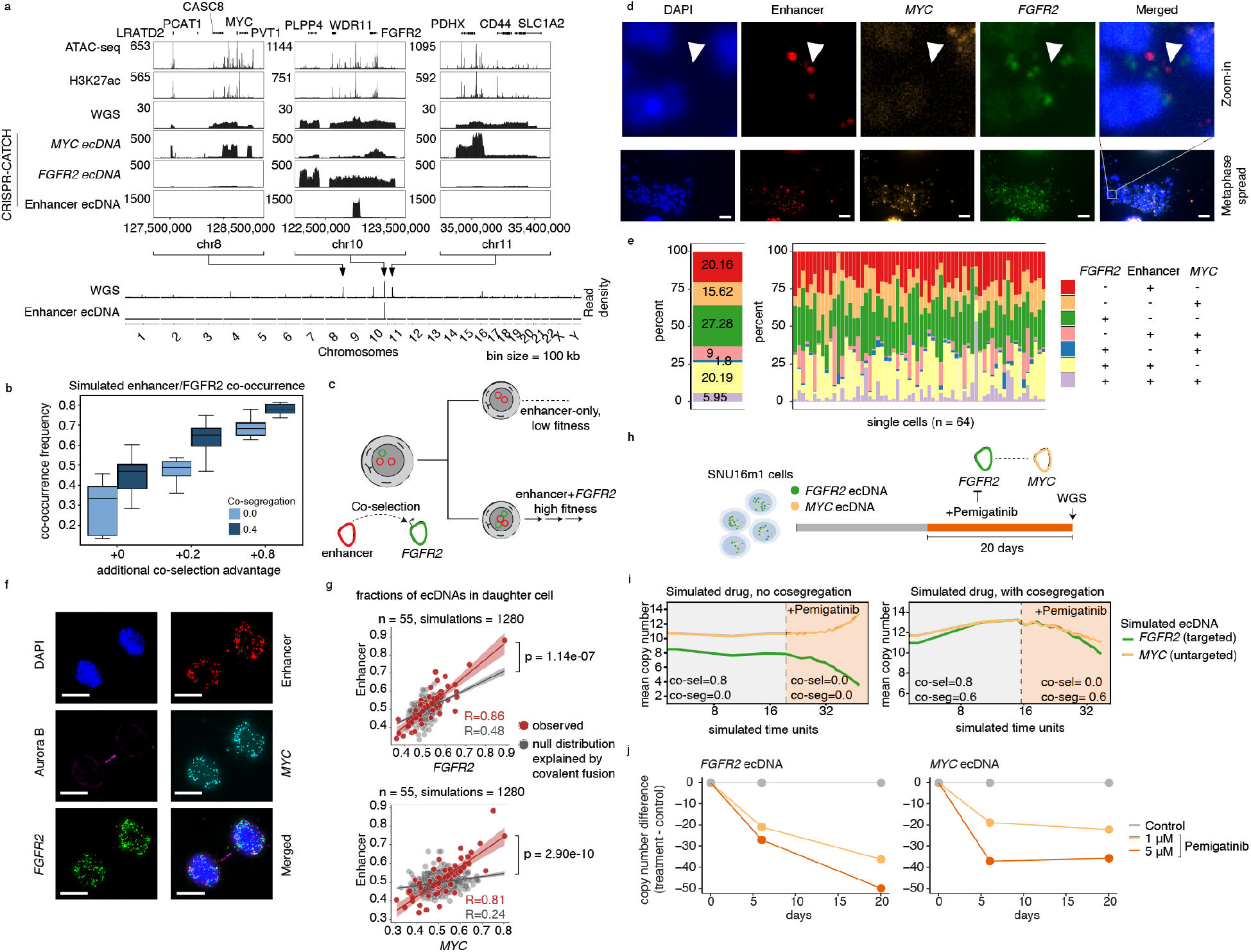
Specialization and therapeutic remodeling of ecDNA species. **(a)** From top to bottom: ATAC-seq, H3K27ac ChIP-seq, WGS, CRISPR-CATCH sequencing of *MYC*, *FGFR2* and enhancer ecDNA species in SNU16 cells; whole-genome read density plots of WGS and CRISPR-CATCH sequencing for enhancer ecDNA. **(b)** Simulated frequencies of cells carrying both *FGFR2* and enhancer-only ecDNA species (i.e., co-occurrence) across varying co-selection and co-segregation values. Co-selection advantage is reported as additive on top of selection acting on cells carrying only *FGFR2* ecDNA. Boxplots show the quartiles of the distribution, and whiskers extend to 1.5x the interquantile range. **(c)** Schematic of co-selection of enhancer ecDNA with *FGFR2* ecDNA. **(d-e)** Representative DNA FISH images of metaphase spreads of SNU16 cells with probes targeting the enhancer, *MYC* and *FGFR2* sequences (d), and quantification of the frequency of each ecDNA species (e). Scale bars are 10 µm. **(f)** Representative images of pairs of SNU16 daughter cells undergoing mitosis identified by immunofluorescence for Aurora kinase B (Aurora B). Individual ecDNA species are visualized using FISH probes targeting specific oncogene or enhancer sequences for copy number quantification. Scale bars are 10 µm. **(g)** Correlation of fractions of ecDNA species in one of each daughter cell pair compared to simulated null distribution explained by covalent fusion (Methods). **(h)** Schematic of *FGFR2* inhibition with Pemigatinib in SNU16m1 cells. **(i)** Computational modeling of *FGFR2* and *MYC* copy numbers in simulated Pemigatinib inhibition experiment with and without co-segregation. Drug efficacy is modeled using decreasing co-selection values. **(j)** Copy-number estimates of *FGFR2* and *MYC* ecDNA in SNU16m1 cells with Pemigatinib treatment or control. Enhancer DNA FISH probe targets the following hg19 genomic coordinates: chr10:123023934-123065872 (WI2-2856M1).

Because co-selection and co-occurrence left distinct signatures on the joint distribution of ecDNA (**Figure 3f**, **Extended Data Figure 4a**), we hypothesized that our evolutionary model could infer levels of ecDNA co-selection and co-segregation from observed single-cell copy number distributions. Pairing our evolutionary model with ecDNA copy number distributions obtained with scATAC-seq, we used Approximate Bayesian Computation (ABC)^24, 25^ to infer posterior distributions for individual selection, co-selection, and co-segregation in the three cell lines containing distinct ecDNA species (**Figure 3g, Extended Data Figure 6a,b**; Methods). As validation, we found that the inferred levels of co-segregation closely matched those observed in paired daughter cells undergoing mitosis using DNA FISH (**Figure 3g**). In line with our previous simulations, we inferred high levels of co-selection of ecDNA species relative to individual selection (**Figure 3f, Extended Data Figure 6b**). Also consistent with our previous simulations, we observed that co-selection coefficients became less critical as we increased the initial copy number for our inference procedure in the population size that we simulated (in effect widening the 95% credible interval for inferred co-selection), while the inferred 95% credible interval of the co-segregation parameter remained stable (**Extended Data Figure 6c**). Together, these results suggest that co-selection and co-segregation underpin the coordinated assortment of ecDNAs in cancer cell populations (**Figure 3h**).

### Behavior of an “altruistic” enhancer-only ecDNA

As our evolutionary model supports the observed distributions and changes in oncogene-encoding ecDNAs, we next asked whether it can also explain the behavior of ecDNAs that do not themselves encode oncogenes but interact with other ecDNA molecules. We recently identified an ecDNA species in the parental SNU16 stomach cancer cell line that did not contain oncogene coding sequences but instead contained a non-coding sequence between *WDR11* and *FGFR2* with accessible chromatin marked by histone H3 lysine 27 acetylation (H3K27ac), suggesting the presence of active enhancers^10^ (**Figure 4a, Extended Data Figure 7a**). These enhancers are required for activation of the *FGFR2* gene and intermolecular activation of the *MYC* gene on ecDNA^9^. Long-read sequencing of the parental cell line revealed that this enhancer ecDNA resulted from two DNA segments joining together by inversions to create a circular molecule (**Extended Data Figure 7a**). As intermolecular interactions of regulatory elements between different ecDNA molecules can drive oncogene expression^9, 17^, the presence of amplified, active enhancer elements in the pool of ecDNA molecules may support enhancer-promoter interactions *in trans* and further upregulate oncogene expression – i.e. an “altruistic” ecDNA. An enhancer-only ecDNA may be especially sensitive to the co-occurrence of oncogene-coding ecDNAs in the same ecDNA hubs to exert its regulatory effect, and presents a unique opportunity to test the predictive framework of our evolutionary model. Simulations under our model of ecDNA co-evolution suggested that co-segregation and co-selection synergize to maintain enhancer-only ecDNAs in a majority of individual cancer cells (**Figure 4b,c**), and that co-selection is particularly important to maintain enhancer-only ecDNAs at substantial copy number in a population (**Extended Data Figure 7b,c)**.

To quantify the frequency of enhancer-only ecDNA species, we performed metaphase DNA FISH with separate, non-overlapping probes targeting *MYC*, *FGFR2* coding sequences, as well as the enhancer sequence, followed by unbiased colocalization analysis of imaging data (Methods). This analysis showed that approximately 20% of ecDNA molecules in SNU16 cells contained this enhancer sequence without either oncogene and that the vast majority of individual cells (98%, 63 of 64 cells examined) harbored the enhancer-only ecDNA species (**Figure 4d,e**). Analysis of pairs of daughter cells undergoing mitosis further showed co-segregation of the enhancer sequence with both *MYC* and *FGFR2* ecDNA molecules significantly above levels that can be explained by covalent linkages alone (*R* > 0.80, *p* < 1e10^-6^ for each comparison, **Figure 4f,g**; Methods). These results support the theory that specialized ecDNAs without oncogenes can arise and be stably maintained by virtue of synergistic interaction with oncogene-carrying ecDNA.

### Pharmacological co-regulation of two interacting ecDNA species

ecDNAs can drive rapid genome evolution in response to pharmacological treatment, including through modulation of ecDNA copy number^12^ and generation of new ecDNAs containing resistance-promoting genes^10, 11^. We hypothesized that co-segregation and co-selection of separate ecDNA species that interact *in trans* could also couple changes in copy number of both ecDNA species in response to targeted drug treatment. Simulations of interventions targeting a single ecDNA species under our evolutionary model predicted a continuous, dose-dependent decrease in copy numbers of co-existing ecDNAs only in the presence of co-segregation (**Figure 4h,i, Extended Data Figure 8a**; Methods). We previously showed that the *MYC* and *FGFR2* ecDNAs engage in intermolecular enhancer-promoter interactions in the SNU16m1 stomach cancer cell line^9^, providing a model in which to test the hypothesis that their copy numbers may be coordinately modulated. We thus hypothesized that treatment of these cells with high dose Pemigatinib, an FGFR2 inhibitor^26, 27^, would remove the selective advantage of cells with amplified *FGFR2* expression and would lead to a loss of *FGFR2* and *MYC* copies over time due to their co-segregating inheritance. This hypothesis is supported by evolutionary modeling (**Figure 4i, Extended Data Figure 8a**). In contrast, in the absence of co-segregation, evolutionary modeling indicates that *FGFR2* and *MYC* ecDNAs would dramatically diverge in their copy numbers, with *MYC* gaining rather than losing copy number upon FGFR2 inhibitor treatment. Experimental treatment of SNU16m1 cells with Pemigatinib over the course of 20 days showed that the *FGFR2* ecDNA copy number decreased progressively in response to drug treatment and in a dose-dependent manner as expected (**Figure 4h,j, Extended Data Figure 8b**). Furthermore, we found that the *MYC* ecDNA species, while not directly targeted by the drug, also decreased in copy number during the course of Pemigatinib treatment (**Figure 4h,j, Extended Data Figure 8b**), supporting the idea that the two ecDNA species are coordinately inherited despite not being covalently linked. Metaphase DNA FISH of drug-treated cells validated the presence of *MYC* and *FGFR2* on separate ecDNAs after Pemigatinib treatment (**Extended Data Figure 8c**), demonstrating that these coordinated decreases in copy number cannot be explained by covalent fusion between ecDNAs. In contrast, Pemigatinib did not result in *MYC* ecDNA loss in the COLO320DM colorectal cancer cell line which does not contain *FGFR2* ecDNAs (**Figure 1c**, **Extended Data Figure 8d**). Consistent with the coordinated change in SNU16m1 cells, targeting *MDM2* ecDNA with Nutlin-3a in TR14 neuroblastoma cells also led to concomitant depletion of co-segregating *MDM2* and *MYCN* ecDNAs, demonstrating the generality of this principle (**Extended Data Figure 8e,f**). Together, these results demonstrate that pharmacological targeting of an oncogene contained on one ecDNA species can coordinately regulate the level of an oncogene on a separate ecDNA species if they co-segregate during mitosis.

## DISCUSSION

Extrachromosomal amplifications in cancer are highly heterogeneous, involving mixtures of species that evolve and increase in complexity over time and in response to selective pressures such as drug treatments^20, 28^. The population of ecDNA species can include amplification of multiple oncogenes, the combination of which provides a fitness advantage to the growing tumor^3, 9–13^. Through single-cell sequencing, DNA FISH, and evolutionary modeling across multiple cancer types, we have shown that diverse ecDNA species co-occur in cancer cells, that they co-segregate with one another during mitosis, and that these evolutionary associations contribute to ecDNA specialization and response to targeted therapy. EcDNAs exhibit aggressive behavior in cancer cells as they can rapidly shift in copy number and evolve novel gene regulatory relationships^4, 9^. This accelerated evolution and ability to explore genetic and epigenetic space is challenged by its potentially transient nature – a winning combination of ecDNAs may not be present in the next daughter cell generation if they are randomly transmitted. EcDNA co-inheritance allows cancer cells to balance accelerated evolution with a measure of genetic and epigenetic memory across cell generations.

While individual ecDNAs are randomly inherited during mitosis^4, 6^, strong co-segregation and co-selection of distinct ecDNAs collaborate to maintain a population of cooperating ecDNAs across generations of cancer cells. Mitotic co-segregation of ecDNAs is imperfect (**Figure 2**); nonetheless, it substantially increases the probability that combinations of ecDNA species will be transmitted together to daughter cells (**Figure 3e**). This coordinated behavior of ecDNA collectives present implications for our understanding of cancer evolution and development of cancer therapies. First, our observation that enhancer-only ecDNAs are co-amplified with *FGFR2* and *MYC* ecDNAs in the SNU16 stomach cancer cells suggests that co-selection of structurally diverse ecDNAs can lead to functional specialization. In light of our recent work describing synergistic intermolecular hubs of ecDNAs^9, 17, 29^, these results suggest that interactive modules of ecDNAs may exist. Second, our findings that ecDNAs can be indirectly depleted through acute inhibition of their co-segregating ecDNA partners (e.g. *FGFR2* and *MYC* ecDNAs or *MDM2* and *MYCN* ecDNAs, respectively) implies that therapeutic interventions targeting the gene product of one ecDNA species can also impact co-amplified oncogenes on other ecDNAs. Importantly, these results do not necessarily imply that oncogene-targeted therapies can “cure” tumor cells of ecDNA, as new ecDNAs carrying resistance-promoting cargo can be selected under targeted therapy. Rather, these results demonstrate that acute targeted therapy can induce rapid, potentially therapeutically advantageous, genomic remodeling as a consequence of ecDNA co-segregation. This indirect targeting of oncogenes may present unique therapeutic opportunities for tumors with co-amplified oncogenes, and we anticipate that the modeling framework described in this study will be a useful resource for understanding when these strategies will be effecitve. Third, our computational model now enables quantification of ecDNA co-segregation and co-selection from genomic or imaging data, including FISH analysis that is widely used in the clinical setting. This advance should enable future research to understand how ecDNAs co-evolve in patient tumors.

The molecular mechanism of ecDNA co-segregation warrants future investigation. As ecDNAs congregate in micron-sized hubs during interphase and interact with one another in an intermolecular manner^9^, one hypothesis is that spatially proximal ecDNAs may co-segregate into daughter cells during mitosis. While ecDNAs lack centromeres and are not attached to the mitotic spindle assembly, they appear to co-localize with mitotic chromosomes and may actively tether to them^5, 9, 30–33^. Many viral episomes tether to mitotic chromosomes during segregation, typically via endogenous and viral protein mediators^34–40^. Non-chromosomal DNA circles in yeast are also retained in the mother cell during mitosis by interacting with protein factors, including nuclear pore complexes^41^. Thus, an alternative and not mutually exclusive hypothesis of the mechanism of ecDNA co-segregation is that ecDNA may rely on a protein or RNA mediator for chromosome tethering and segregation, and asymmetric partitioning of one or more mediators may determine inheritance of multiple ecDNA species by a daughter cell. Future work may search for such mediators by screening for small molecules or genetic perturbations that alter ecDNA co-segregation.

Together, our work identifies the principles of how intermolecular interactions between distinct ecDNAs are preserved amidst asymmetric segregation. The consequence is a jackpot effect that supports cooperation among heterogeneous ecDNAs, enabling the co-selection and co-amplification of multiple oncogenes and continued diversification of cancer genomes. Just as quantitative understanding of chromosome assortment provided the basis of genetic linkage analysis, a quantitative understanding of ecDNA inheritance and deviations from expected patterns may have additional dividends. Beyond cancer evolution, our general framework for coordinated asymmetric inheritance may be applicable to viral episomes, subcellular organelles, or biomolecular condensates that control cell fates.

## Acknowledgements

We thank members of the Chang and Mischel labs for discussion, as well as Jeremy D’Silva for helpful suggestions in developing an evolutionary model of ecDNA co-inheritance. This project was supported by Cancer Grand Challenges CGCSDF-2021 100007 with support from Cancer Research UK and the National Cancer Institute (H.Y.C., V.B., P.S.M., A.G.H.); NIH R35-CA209919 (H.Y.C.); U24CA264379, R01GM114362, OT2CA278635, and CGCATF-2021/100025 (V.B.). K.L.H. was supported by a Stanford Graduate Fellowship and an NCI Predoctoral to Postdoctoral Fellow Transition Award (NIH F99CA274692). J.A.B. was supported by NIH training grant T32HL120824. A.G.H. is supported by the Deutsche Forschungsgemeinschaft (DFG, German Research Foundation) – 398299703 and the European Research Council (ERC) under the European Union’s Horizon 2020 research and innovation programme (grant agreement No. 949172). B.W. is supported by a Barts Charity Lectureship (MGU045) and a UKRI Future Leaders Fellowship (MR/V02342X/1). H.Y.C. is an Investigator of the Howard Hughes Medical Institute.

## Author Contributions

K.L.H., J.T.L., P.M. and H.Y.C. conceived the project. K.L.H. analyzed single-cell ATAC-seq and RNA-seq data, analyzed ecDNA copy number and colocalization using metaphase DNA FISH images, analyzed ecDNA segregation in mitotic immunofluorescence and DNA FISH images, performed simulations of ecDNA segregation in paired daughter cells, performed CRISPR-CATCH experiments and analyses, integrated ATAC-seq and ChIP-seq data, and analyzed WGS data. M.G.J. performed evolutionary modeling, conducted Pemigatinib treatments in cell culture and performed Nanopore sequencing of SNU16 genomic DNA. I.T.L.W. and J.T.L. performed immunofluorescence staining and DNA FISH in mitotic cells and imaging. J.L. analyzed ecDNA amplicon sequences in TCGA patient tumors using AmpliconClassifier. E.S. created the alternative model of ecDNA co-evolution. K.L.H., B.J.H. and R.L. prepared sequencing libraries for WGS and CRISPR-CATCH. K.L.H. and R.L. prepared sequencing libraries for single-cell ATAC-seq and RNA-seq. I.T.L.W., L.B., R.S., J.R.D. performed DNA FISH and imaging experiments. R.C.G. analyzed scCircle-seq data. J.A.B., B.W., W.H., V.B., A.G.H., P.S.M. and H.Y.C. guided data analysis and provided feedback on experimental design. K.L.H., M.G.J. and H.Y.C. wrote the manuscript with input from all authors.

## Extended Data Figures

**Extended Data Figure 1.**
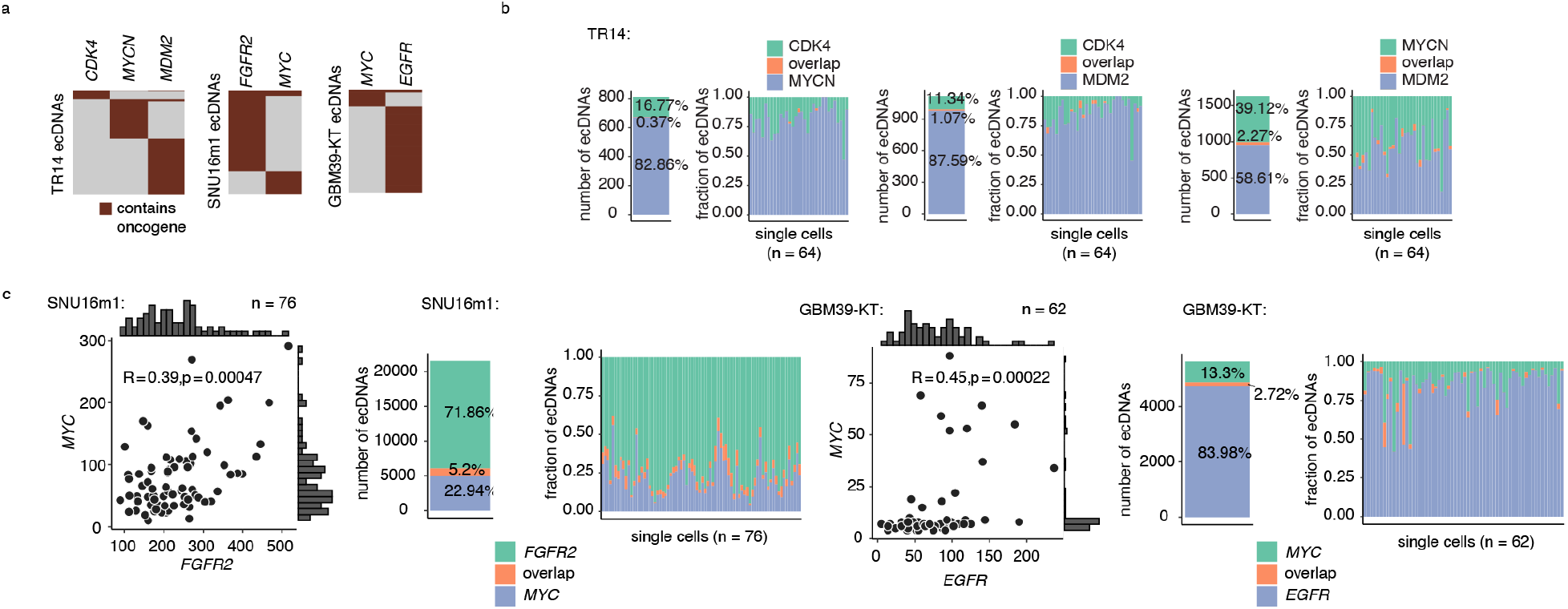
Oncogenes are harbored on distinct ecDNA species but are correlated in copy number. **(a)** Heatmaps showing non-overlapping oncogene presence on distinct ecDNA species in metaphase DNA FISH. Rows represents individual ecDNA molecules. **(b)** Bar plots showing the fractions of ecDNAs containing combinations of MYCN, CDK4 or MDM2 and demonstrating little overlap between these oncogenes on the same ecDNA molecules. **(c)** Copy number correlations and distributions of oncogene ecDNAs in metaphase DNA FISH images (left), and bar plots showing the fractions of ecDNAs containing combinations of *FGFR2* and *MYC* in SNU16m1 cells and combinations of *MYC* and *EGFR* in GBM39-KT cells, demonstrating little overlap between these oncogenes on the same ecDNA molecules (right).

**Extended Data Figure 2.**
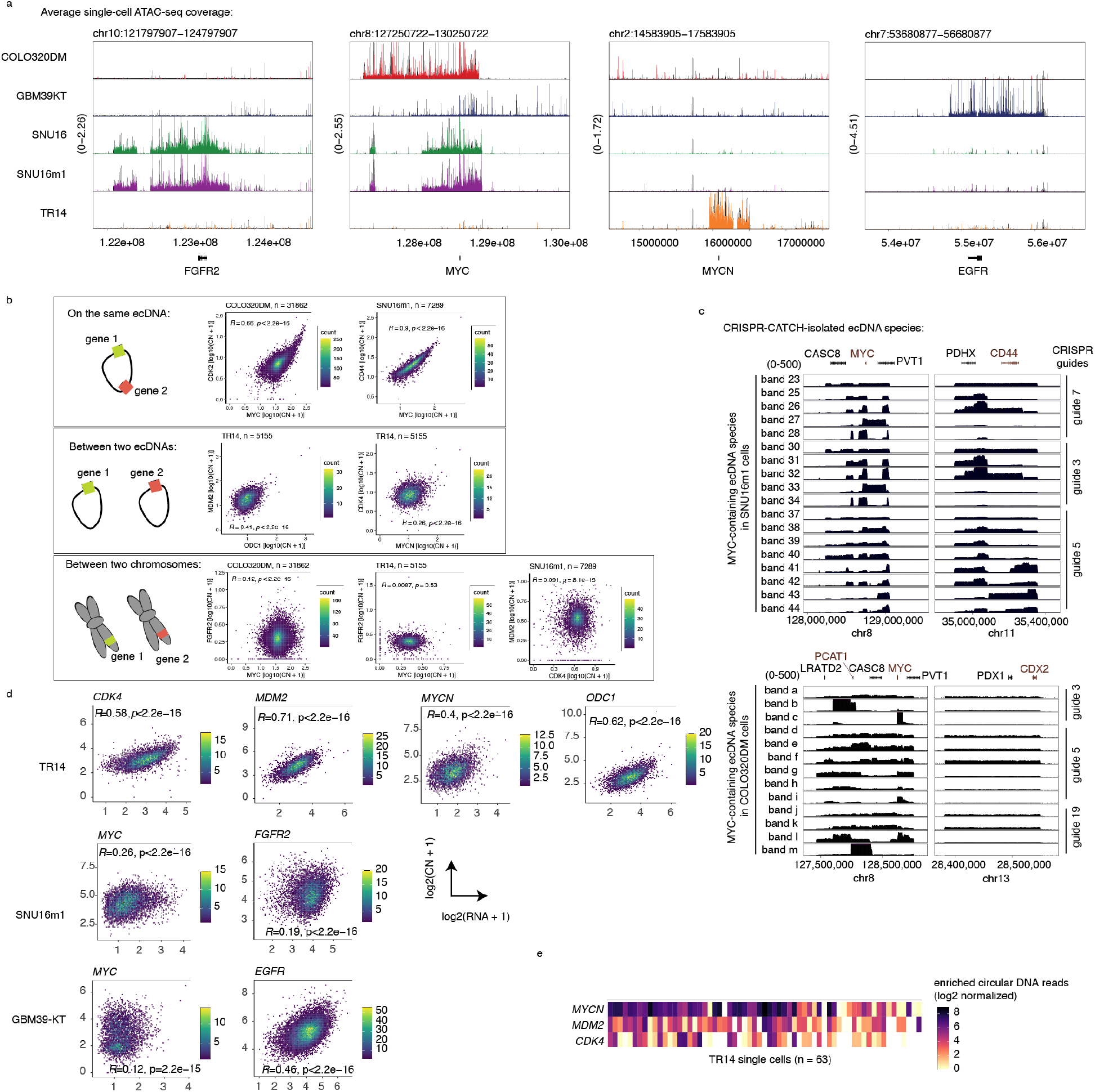
Distinct ecDNA amplifications co-occur and correlate at the single-cell level and their copy numbers affect transcriptional outcomes of oncogenes. **(a)** Elevated scATAC-seq background coverages of oncogene loci in correspondence to ecDNA copy number amplification in the various indicated cell lines. **(b)** Density scatter plots showing levels of copy number correlation between gene pairs on the same ecDNA, on different ecDNAs, and on different chromosomes. **(c)** Sequencing coverages of ecDNA species isolated by CRISPR-CATCH from SNU16m1 cells and COLO320DM cells, identifying genes that are frequently linked on the same ecDNA species **(Methods)**. Each row represents a distinct ecDNA species isolated by molecular size fractionation using CRISPR-CATCH. Gene annotations in red are gene pairs classified as being on the same ecDNA in Figure 1. All guide sequences are provided in Supplementary Table 1. **(d)** Density scatter plots showing correlation between oncogene copy number and RNA expression in paired scATAC-seq and RNA-seq. Cells with zero values were filtered. **(e)** Heatmap showing co-enrichment of circular DNA species containing *MYCN*, *MDM2* or *CDK4* in individual TR14 neuroblastoma cells in scCircle-seq.

**Extended Data Figure 3.**
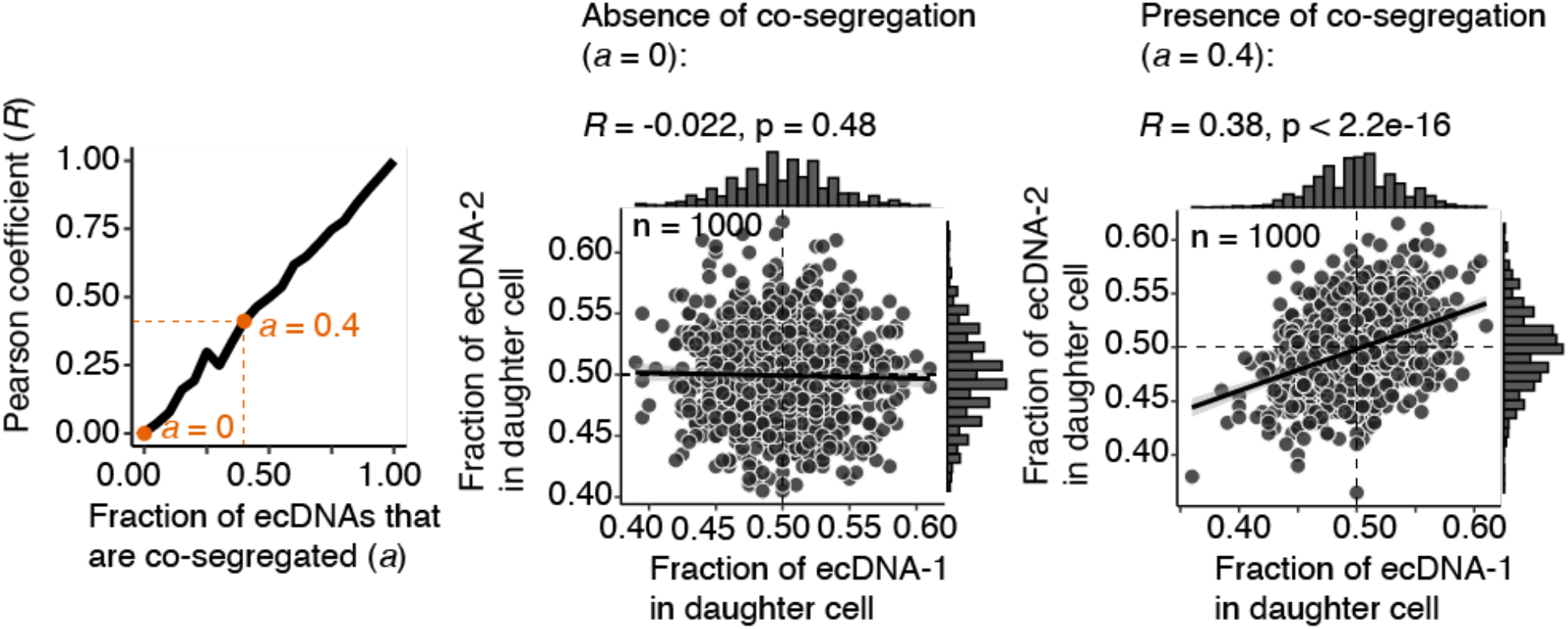
Copy number correlation between ecDNA species in daughter cells undergoing mitosis suggests co-segregation. Segregation of two ecDNA species with 100 copies each was simulated by random sampling with varying levels of co-segregation (1000 simulations per co-segregation fraction *a*; Methods). As the fraction (*a*) of ecDNAs that are co-segregated increases from 0.00 (no co-segregation) to 1.00 (each copy of one ecDNA species is perfectly co-segregated with a copy of another species) in increments of 0.05, the Pearson coefficient *R* of the copy numbers of two ecDNA species in individual daughter cells increases linearly (left panel). Thus, in the absence of co-segregation, no copy number correlation in mitotic daughter cells is expected (middle panel), while in the presence of a modest level of co-segregation (a fraction of 0.4, or 40% of one ecDNA species co-segregating with 40% of another), a Pearson coefficient *R* of 0.38 is expected (right panel).

**Extended Data Figure 4.**
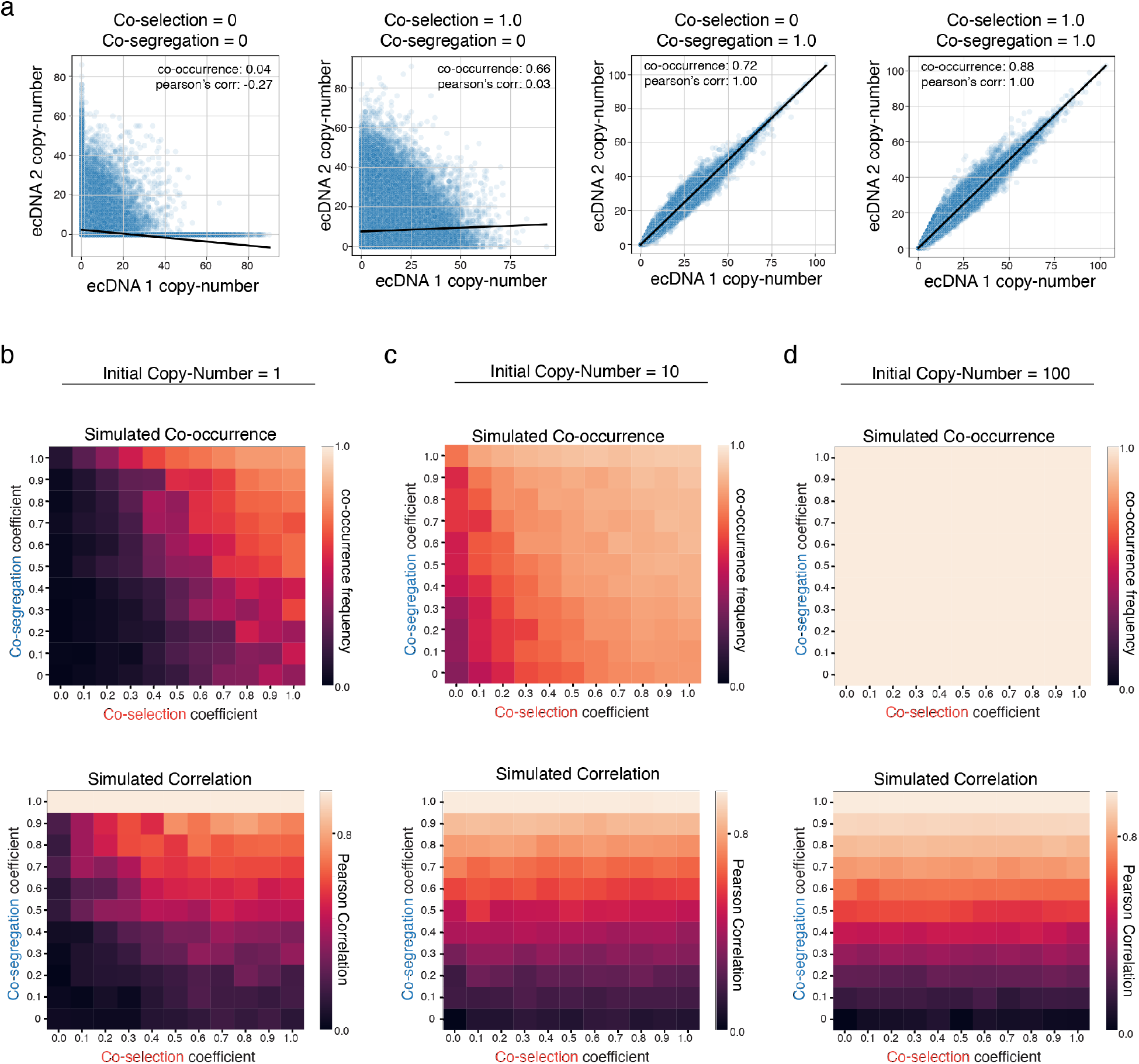
Characterization of ecDNA evoluationary model across various initial conditions. **(a)** Representative joint ecDNA copy-number distributions across varying levels of co-segregation and co-selection. Co-occurrence frequency and Pearson’s correlation are reported for each joint distribution. **(b-d)** Average frequency of cells carrying both ecDNA species and Pearson’s correlation of ecDNA copy-numbers in single cells are reported across simulations of 10 replicates of 1 million cells for varying initial ecDNA copy-numbers: **(b)** 1 copy of each ecDNA species; **(c)** 10 copies of each ecDNA species; **(d)** 100 copies of each ecDNA species. Selection acting on cells with only one but not both ecDNA species is maintained at 0.2 and selection acting on cells without either ecDNA is maintained at 0.0 for all simulations.

**Extended Data Figure 5.**
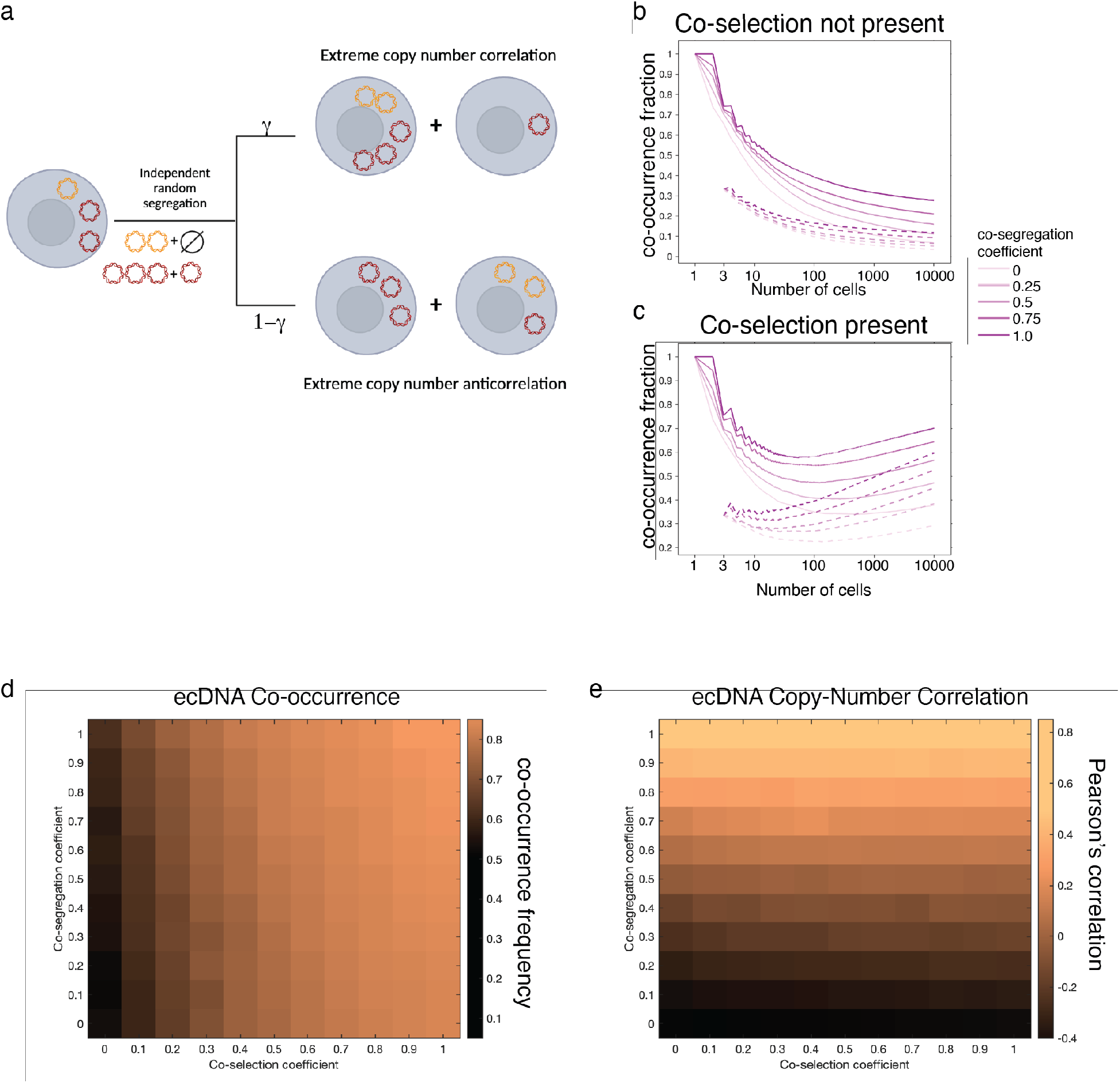
EcDNA co-inheritance dynamics using an alternative model of ecDNA evolution. **(a)** A schematic illustrating an alternative model of ecDNA evolution, paramterized by selection acting on cells carrying no, both, or either ecDNA as well as a co-segregation parameter *γ*. **(b-c)** Frequency of cells carrying both ecDNA species reported as a function of number of cells during a simulation for variable levels of co-segregation and with **(b)** or without **(c)** co-selection. **(d-e)** Average frequencies of cells carrying both ecDNA species **(d)** and the Pearson’s correlation of ecDNA copy numbers across 500 replicates of simulations of 10,000 cells while varying co-selection and co-segregation values.

**Extended Data Figure 6.**
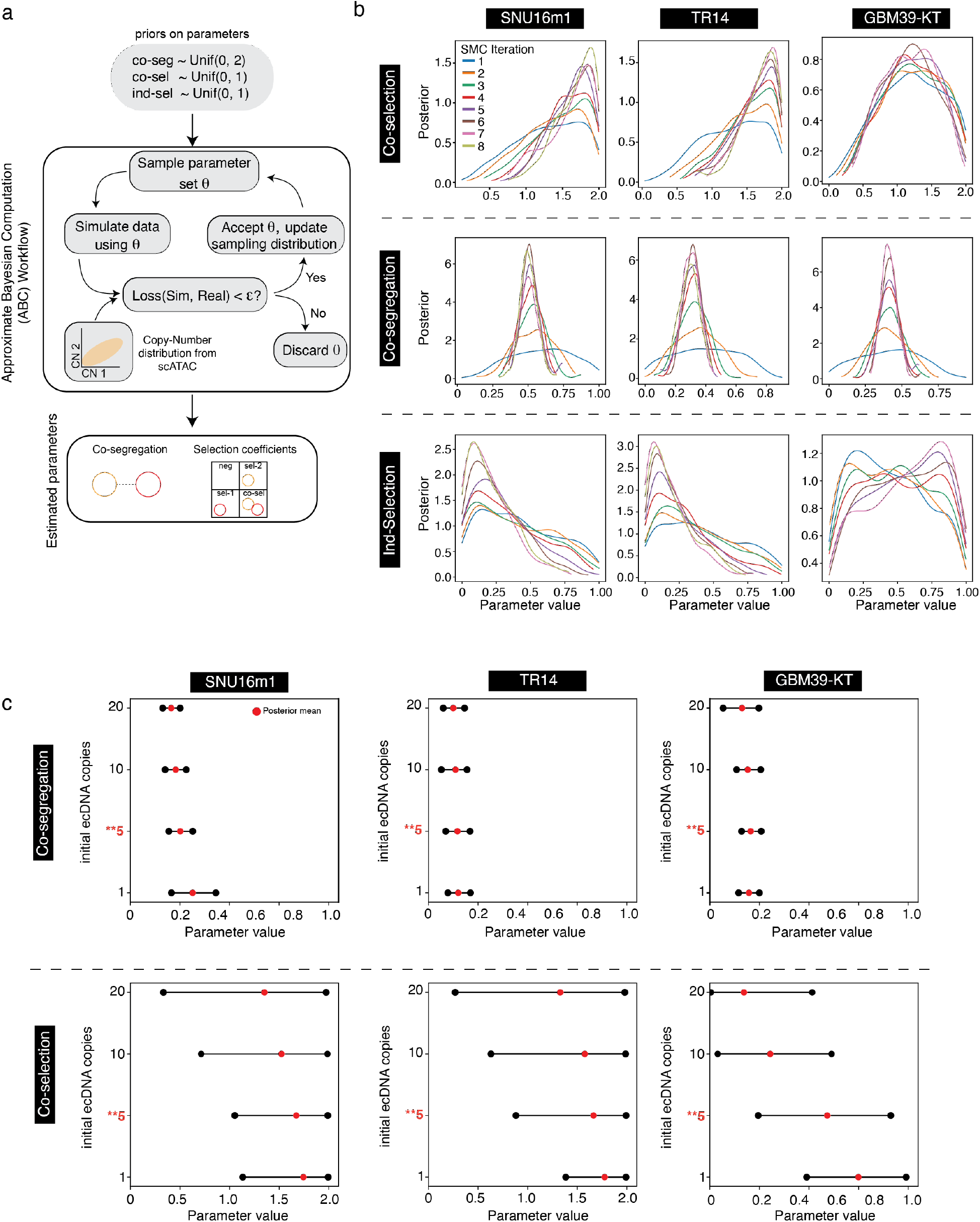
Analysis of Approximate Bayesian Computation (ABC) inference stability. **(a)** Schematic of ABC inference workflow: posterior distributions over parameters are inferred from user-defined priors and observed single-cell copy-number data using sequential model fitting on our evoluationy model. **(b)** Posterior distributions of co-selection, co-segregation, and individual selection values for inferences in SNU16m1, TR14, and GBM39-KT across sequential iterations of Approximate Bayesian Inference Sequential Monte Carlo (ABC-SMC). **(c)** 95% credible interval of inferred co-segregation and co-selection values from ABC-SMC across the cell lines studied in this report with variable initial ecDNA copy numbers (1, 5, 10, 20). The initial ecDNA copy number (5) used in the main text is highlighted in red.

**Extended Data Figure 7.**
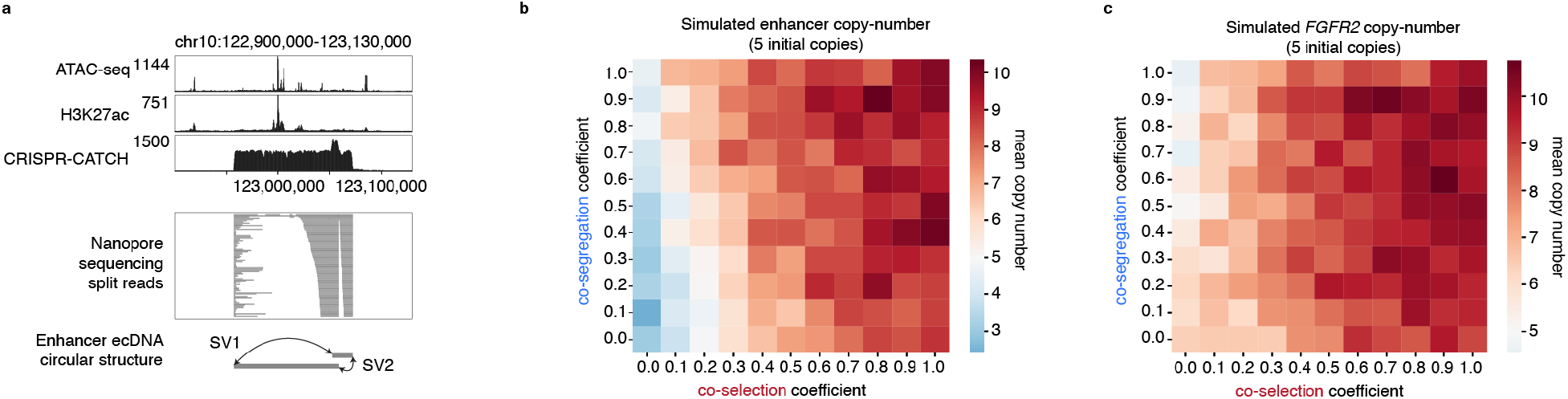
Structure and dynamics of the enhancer ecDNA. **(a)** From top to bottom: ATAC-seq, H3K27ac ChIP-seq, CRISPR-CATCH sequencing of enhancer-only ecDNA species in SNU16 cells, individual split reads in Nanopore sequencing supporting the circular enhancer-only ecDNA species, and structural variants (SV1 and SV2) that create a circular structure. SV1: precise inversion between chr10:122957191 and chr10:123051954; SV2: precise inversion between chr10:123058196 and chr10:123071737. **(b-c)** Simulated copy number of enhancer-only ecDNA (**b**) and *FGFR2* ecDNA (**c**) under various settings of co-selection and co-segregation. Individual selection on the enhancer-only species was kept at 0.0, and individual selection on the *FGFR2* ecDNA was kept at 0.2. One million cells were simulated from a parent cell carrying 5 copies of both species. 10 replicates were simulated and the average value was reported.

**Extended Data Figure 8.**
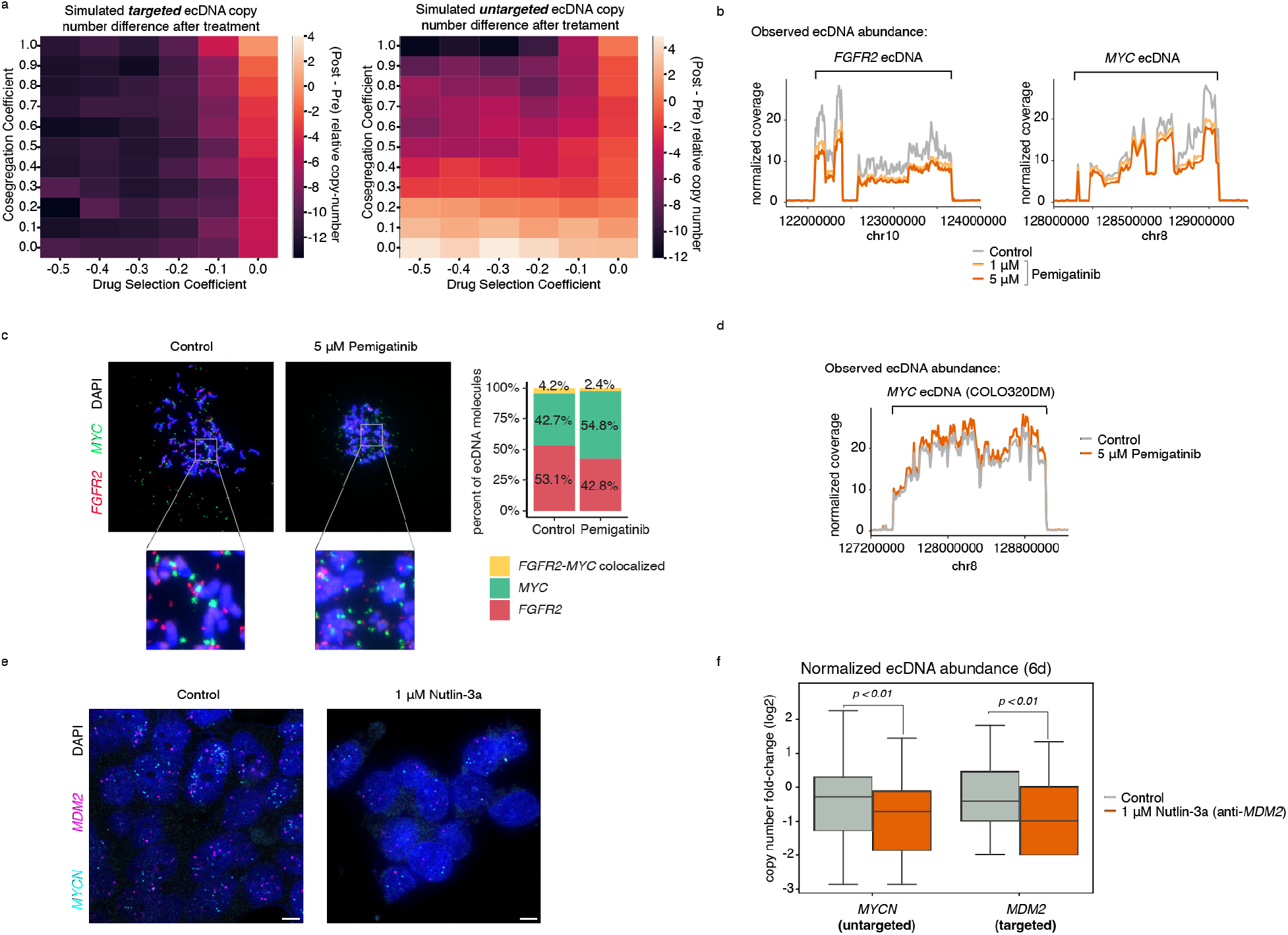
Pemigatinib results in coordinated decreases in ecDNA copy numbers but not via oncogene fusion or direct targeting of MYC. (**a**) Simulated changes in copy number after targeted treatment for the ecDNA directly or indirectly being targeted under various parameters of co-segregation and drug selection. 500,000 cells were simulated, and average values were reported across 10 replicates. **(b)** WGS coverage of *FGFR2* and *MYC* ecDNA genomic intervals after 20 days of Pemigatinib treatment at 1 *μ*M and 5 *μ*M compared to PBS control. (**c)** Representative metaphase DNA FISH images showing distinct *FGFR2* and *MYC* ecDNA species in SNU16m1 cells after 20 days of treatment with 5 *μ*M Pemigatinib or PBS control (left), and quantification of distinct and colocalized *FGFR2-MYC* DNA FISH signals (right). **(d)** WGS coverage of *MYC* ecDNA genomic interval in COLO320DM cells after 20 days of treatment with 5 μM Pemigatinib compared to PBS control. **(e)** Representative images of DNA FISH on interphase TR14 cells with and without 1 *μ*M Nutlin-3a treatment after 6 days. Scale bars are 5 µm. **(f)** Normalized copy number of *MDM2* and *MYCN* in TR14 cells after 6 days of 1 *μ*M Nutlin-3a or DMSO control treatment (p-values computed with a one-sided Wilcoxon rank-sums test). Boxplots show the quartiles of the distribution, and whiskers extend to 1.5x the interquantile range.

## METHODS

### Cell culture

The TR14 neuroblastoma cell line was a gift from J. J. Molenaar (Princess Máxima Center for Pediatric Oncology, Utrecht, Netherlands). Cell line identity for the master stock was verified by STR genotyping (IDEXX BioResearch, Westbrook, ME). The GBM39-KT cell line was derived from a patient with glioblastoma as described previously^7^. Parental SNU16 was obtained from ATCC. The monoclonal SNU16m1 was a subline of the parental SNU16 cells generated from a single cell after lentiviral transduction and stable expression of dCas9-KRAB as we previously described^9^. SNU16 and SNU16m1 cells were maintained in Dulbecco’s modified Eagle’s medium/nutrient mixture F-12 (DMEM/F12 1:1; Gibco, 11320-082), 10% fetal bovine serum (FBS; Hyclone, SH30396.03) and 1% pen-strep (Thermo Fisher Scientific, 15140-122). COLO320-DM cells were maintained in DMEM (Thermo Fisher Scientific, 11995073) supplemented with 10% FBS and 1% pen-strep. GBM39-KT cells were maintained in DMEM/F12 1:1, B-27 supplement (Gibco, 17504044), 1% pen-strep, GlutaMAX (Gibco, 35050061), human epidermal growth factor (EGF, 20 ng ml−1; Sigma-Aldrich, E9644), human fibroblast growth factor (FGF, 20 ng ml−1; Peprotech) and heparin (5 μg ml−1; Sigma-Aldrich, H3149-500KU). TR14 cells were grown in RPMI 1640 with 20% FBS and 1% pen-strep. For the mitotic cell imaging experiments in Figure 2, SNU16m1 cells were grown in Roswell Park Memorial Institute (RPMI) 1640 with 10% FBS. All cells were cultured at 37 °C with 5% CO2. All cell lines tested negative for mycoplasma contamination.

### WGS

WGS libraries were prepared by DNA tagmentation. We first transposed it with Tn5transposase produced as previously described^42^, in a 50-µl reaction with TD buffer^43^, 50 ng DNA and 1 µl transposase. The reaction was performed at 50°C for 5 minutes, and transposed DNA was purified using MinElute PCR Purification Kit (Qiagen, 28006). Libraries were generated by 5-7 rounds of PCR amplification using NEBNext High-Fidelity 2× PCR Master Mix (NEB, M0541L), purified using SPRIselect reagent kit (Beckman Coulter, B23317) with double size selection (0.8× right, 1.2× left) and sequenced on the Illumina Nextseq 550 or the Illumina NovaSeq 6000 platform. Reads were trimmed of adapter content with Trimmomatic^44^ (version 0.39), aligned to the hg19 genome using BWA MEM^45^ (0.7.17-r1188), and PCR duplicates removed using Picard’s MarkDuplicates (version 2.25.3). WGS data from bulk SNU16 cells were previously generated (SRR530826, Genome Research Foundation).

### Analysis of ecDNA sequences in TCGA patient tumors

We utilized AmpliconArchitect (v1.0) outputs from Kim et al. 2020^2^, and classified focal amplifications types present in these outputs using AmpliconClassifier (v0.4.14) with the “—filter_similar” flag set and otherwise default settings. The “—filter_similar” option removes likely false positive focal amplification calls which contain far greater-than-expected levels of overlapping SVs and shared genomic boundaries between ecDNAs of unrelated samples. Of 8810 AA amplicons in the Kim et al. TCGA dataset, 45 were removed by this filter. Individual samples were considered to have a number of ecDNA species equal to the total number of ecDNA species predicted in each AA amplicon, across all AA amplicons detected in the sample. A list of oncogenes was created using genes in the the ONGene database (https://pubmed.ncbi.nlm.nih.gov/28162959/) and COSMIC (https://www.ncbi.nlm.nih.gov/pmc/articles/PMC6450507/).

### Paired single-cell ATAC-seq and RNA-seq library generation

Single-cell paired RNA and ATAC-seq libraries were generated on the 10x Chromium Single-Cell Multiome ATAC + Gene Expression platform following the manufacturer’s protocol and sequenced on an Illumina NovaSeq 6000. Data for COLO320DM were generated previously in Hung et al. 2021^9^ and published under GEO accession GSE159986.

### Paired single-cell ATAC-seq and RNA-seq analysis

A custom reference package for hg19 was created using cellranger-arc mkref (10x Genomics, version 1.0.0). The single-cell paired RNA and ATAC-seq reads were aligned to the hg19 reference genome using cellranger-arc count (10x Genomics, version 1.0.0).

Subsequent analyses on RNA were performed using Seurat (version 3.2.3)^46^, and those on ATAC-seq were performed using ArchR (version 1.0.1)^47^. Cells with more than 200 unique RNA features, less than 20% mitochondrial RNA reads, less than 50,000 total RNA reads were retained for further analyses. Doublets were removed using ArchR. Raw RNA counts were log-normalized using Seurat’s NormalizeData function and scaled using the ScaleData function. Dimensionality reduction for the ATAC-seq data were performed using Iterative Latent Semantic Indexing (LSI) with the addIterativeLSI function in ArchR. We then calculated amplicon copy numbers based on background ATAC-seq signals as we previously described and validated^9, 48^. Briefly, we determined read counts in large intervals across the genome using a sliding window of three megabases moving in one-megabase increments across the reference genome. Genomic regions with known mapping artifacts were filtered out using the ENCODE hg19 blacklist. For each interval, insertions per base pair were calculated and compared to 100 of its nearest neighbors with matched GC nucleotide content. Mean log_2_(fold change) was computed for each interval. Based on a diploid genome, copy numbers were calculated using the formula CN = 2 * [2 ∧ log_2_(FC)], where CN denotes copy number and FC denotes mean fold change compared to neighboring intervals. To query the copy numbers of a gene, we obtained all genomic intervals that overlapped with the annotated gene sequence and and computed the mean copy number of those intervals.

### Single-cell Circle-seq

TR14 scCircle-seq data were deposited in the European Genome-phenome Archive (EGA) under the accession number: EGAS00001007026. A detailed description of the single-cell extrachromosomal circular DNA and transcriptome sequencing (scEC&T-seq) protocol is available on *Nature protocol exchange* (DOI: 10.21203/rs.3.pex-2180/v1). In short, single cells were separated into individual wells of a 96-well plates using FACS. Separation of genomic DNA and mRNA was performed as described in the G&T-seq protocol by Macaulay et al. 2015^49^. Genomic DNA of single cells was purified using 0.8× AMPure XP beads and subjected to exonuclease digestion and rolling-circle amplification as described in Chamorro González et al, 2023 (in press). All single-cell libraries were prepared using the NEBNext Ultra II FS kit (New England Biolabs) following the manufacturer’s instructions but using one-fourth volumes. Unique dual index primer pairs (New England Biolabs) were used to barcode single-cell libraries. Pooled libraries were sequenced on the Illumina HiSeq 4000 or the NovaSeq 6000 platform with 2 × 150bp paired-end reads for genomic DNA and circular DNA libraries and 2 × 75 bp paired-end reads for cDNA libraries.

### ecDNA isolation by CRISPR-CATCH

Molecular isolation of ecDNA by CRISPR-CATCH was performed as previously described^10^. Briefly, molten 1% certified low melt agarose (Bio-Rad, 1613112) in PBS was equilibrated to 45°C. 1 million cells were pelleted per condition, washed twice with cold 1× PBS, resuspended in 30 µl PBS, and briefly heated to 37°C. 30 µl agarose solution was added to cells, mixed, transferred to a plug mold (Bio-Rad Laboratories, Cat #1703713) and incubated on ice for 10 minutes. Solid agarose plugs containing cells were ejected into 1.5-ml Eppendorf tubes, suspended in buffer SDE (1% SDS, 25 mM EDTA at pH 8.0) and placed on a shaker for 10 minutes. The buffer was removed and buffer ES (1% N-laurolsarcosine sodium salt solution, 25 mM EDTA at pH 8.0, 50 µg/ml proteinase K) was added. Agarose plugs were incubated in buffer ES at 50°C overnight. On the following day, proteinase K was inactivated with 25 mM EDTA with 1 mM PMSF for 1 hour at room temperature with shaking. Plugs were then treated with RNase A (1 mg/ml) in 25 mM EDTA for 30 minutes at 37°C, and washed with 25 mM EDTA with a 5-minute incubation. Plugs not directly used for ecDNA enrichment were stored in 25 mM EDTA at 4°C.

To perform *in-vitro* Cas9 digestion, agarose plugs containing DNA were washed three times with 1× NEBuffer 3.1 (New England BioLabs) with 5-minute incubations. Next, DNA was digested in a reaction with 30 nM single-guide RNA (sgRNA, Synthego) and 30 nM spCas9 (New England BioLabs, M0386S) after pre-incubation of the reaction mix at room temperature for 10 minutes. Cas9 digestion was performed at 37°C for 4 hours, followed by overnight digestion with 3 µl proteinase K (20 mg/ml) in a 200 µl reaction. On the following day, proteinase K was inactivated with 1 mM PMSF for 1 hour with shaking. plugs were then washed with 0.5× TAE buffer three times with 5-minute incubations. Plugs were loaded into a 1% certified low melt agarose gel (Bio-Rad, 1613112) in 0.5× TAE buffer with ladders (CHEF DNA Size Marker, 0.2–2.2 Mb, *Saccharomyces cerevisiae* Ladder: Bio-Rad, 1703605; CHEF DNA Size Marker, 1–3.1 Mb, *Hansenula wingei* Ladder: Bio-Rad, 1703667) and PFGE was performed using the CHEF Mapper XA System (Bio-Rad) according to the manufacturer’s instructions and using the following settings: 0.5× TAE running buffer, 14°C, two-state mode, run time duration of 16 hours 39 minutes, initial switch time of 20.16 seconds, final switch time of 2 minutes 55.12 seconds, gradient of 6 V/cm, included angle of 120°, and linear ramping. Gel was stained with 3× Gelred (Biotium) with 0.1 M NaCl on a rocker for 30 minutes covered from light and imaged. Bands were then extracted and DNA was isolated from agarose blocks using beta-Agarase I (New England BioLabs, M0392L) following the manufacturer’s instructions. All guide sequences are provided in **Supplementary Table 1**.

### Short-read sequencing of ecDNA isolated by CRISPR-CATCH

Sequencing of ecDNA isolated by CRISPR-CATCH was performed as previously described^10^. Briefly, we transposed DNA with Tn5 transposase produced as previously described^42^, in a 50-µl reaction with TD buffer^43^, 10 ng DNA and 1 µl transposase. The reaction was performed at 50°C for 5 minutes, and transposed DNA was purified using MinElute PCR Purification Kit (Qiagen, 28006). Libraries were generated by 7-9 rounds of PCR amplification using NEBNext High-Fidelity 2× PCR Master Mix (NEB, M0541L), purified using SPRIselect reagent kit (Beckman Coulter, B23317) with double size selection (0.8× right, 1.2× left) and sequenced on the Illumina Nextseq 550 or the Illumina NovaSeq 6000 platform. Sequencing data were processed as described above for WGS. CRISPR-CATCH sequencing data for SNU16m1 (bands 29-34) and COLO320DM (bands a-m) used in **Extended Data Figure 1** were generated previously in Hung et al. 2021^9^ and deposited in NCBI Sequence Read Archive (SRA) under BioProject accession PRJNA670737; CRISPR-CATCH sequencing data for SNU16 (*MYC*, *FGFR2* and enhancer ecDNAs) used in **Figure 4** were generated previously in Hung et al. 2022^10^ and deposited in NCBI SRA under BioProject accession PRJNA777710.

### Metaphase DNA FISH

TR14 neuroblastoma cells were grown to 70% confluency in a 15 cm dish and treated with KaryoMAX™ Colcemid™ (Gibco) for four hours. A mitotic shake off was preformed and the media of the cells was collected. The remaining cells were washed with PBS and treated with Trypsin-EDTA 0,05 % (Gibco) for 2 minutes. The cells were washed again with the collected media and spun down at 300 g for 10 minutes. The pellet was resuspended at 0.075M KCl and left at 37°C for 20 minutes. The sample was spun down at 300 g for 5 minutes. The cell pellet was resuspended carefully in 10 mL Carnoy’s solution and spun down at 300 g for 5 minutes. This washing step was repeated 4 times using 5 mL of Carnoy’s solution. The remaining pellet was resuspended in 400 µl of Carnoy’s solution. 12 µl of the suspension was dropped on preheated slides from a height of approximately 15 cm. The slides were held over a heated water bath (55 °C) for one minute. Slides were aged overnight at room temperature. Slides were prepared for staining following the probe manufacturer’s protocol (DNA FISH Metaphase Chromsome spreads, Arbor Biosciences). Before staining, slides were firstly washed in PBS, followed by a wash in 65°C SSCT (5 mL 20x SSC, 500 µl 10% Tween 20, bring up to 50 mL with molecular grade H20) for 15 minutes. Afterwards, slides were washed 2 times for 2 minutes with room temperature SSCT. Dehydration of the slides was done in 70% and 90% ethanol for 5 minutes each. After airdrying, slides were transferred into 0.07 N NaOH for 3 minutes for chemical denaturation. After 2 washes for 5 minutes in SSCT, the dehydration step was repeated, and slides were airdried. The probes used for staining were designed to target the *MYCN, MDM2* and *CDK4* gene by myTags^®^ (Arbor), conjugated as following: CDK4-Alexa 488, MYCN-Atto 550, MDM2-Atto 633. 10 µl of the hybridization buffer (in SSCT: 50 % formamide, 10% dextran sulphate, 40 ng/µl RNase A) were mixed with 1,5 µl of each resuspended probe. This mixture was headed to 70 °C for 5 minutes and stored on ice. 14,5 µl were added to the slide, covered by a cover glass, and sealed with rubber cement. The slides were incubated in a hybridization chamber (Abbott Molecular) overnight at 37 °C. On the next day, rubber cement and cover glass were removed, and the sample was washed in prewarmed (37 °C) SSCT for 30 minutes. Afterwards, slides were washed at room temperature with 2 times SSCT for 5 minutes each followed by a 5 minutes wash with PBS. The air-dried slide was stained with Hoechst (1: 4000 for 2 minutes) and washed with PBS for another 5 minutes. After drying, the slides were mounted using ProLong Glass Antifade Mountant (ThermoFisher Scientific) and sealed with a coverglass. Imaging of TR14 metaphase spreads was done on the Leica Stellaris 8 (Advanced light microscopy facility, Max-Delbrück Center for molecular medicine) using a 63x oil objective with a 2x zoom. Excitation was done using the 405, 488, 561 and 538 lasers and detection was done using two HyD S and one HyD X and HyD R detectors. 4x line averaging was applied to each channel.

For the GBM39-KT, SNU16 and SNU16m1 cell ines, cells were treated with KaryoMAX Colcemid (Gibco) at 100 ng ml−1 for 3 hours, and single cell suspensions were then collected by centrifugation and washed once in 1× PBS. The cells were treated with 0.75M KCl hypotonic buffer for 20 minutes at 37 °C, and fixed with Carnoy’s fixative (3:1 methanol:glacial acetic acid) followed by three additional washes with the same fixative. The samples were then dropped onto humidified glass slides and air-dried. The glass slides were then briefly equilibrated in 2× SSC buffer, dehydrated in ascending ethanol concentrations of 70%, 85% and 100% for 2 minutes each. FISH probes (Empire Genomics) were diluted in hybridization buffer in 1:6 ratio and covered with a coverslip. Samples were denatured at 75 °C for 3 minutes and hybridized at 37 °C overnight in a humidified slide moat. The samples were washed with 0.4× SSC for 2 minutes, and 2× SSC 0.1% Tween 20 for another 2 minutes. The nuclei were stained with 4,6-Diamidino-2-phenylindole (DAPI) (50 ng ml−1) diluted in 2× SSC for about a minute, and washed once briefly in ddH2O. Air-dried samples were mounted with ProLong Diamond. Images were acquired on a Leica DMi8 widefield microscope using a 63× oil objective.

### Metaphase DNA FISH image analysis

Colocalization analysis for two- and three-color metaphase FISH described in **Figure 1** and **Extended Data Figure 1** was performed using Fiji (version 2.1.0/1.53c)^50^. Images were split into the individual FISH colors + DAPI channels, and signal threshold set manually to remove background fluorescence. Overlapping FISH signals were segmented using watershed segmentation. FISH signals were counted using particle analysis. XY coordinates of pixels containing FISH signals were saved along with image dimensions and coordinates of regions of interest (ROIs) as distinct particle identities (e.g. distinct ecDNA molecules). Colocalization was then quantified in R. Each pixel containing FISH signal was assigned to the nearest overlapping ROI using XY coordinates. Unique ROIs in all color channels were summarized such that ROIs in different channels that overlap with one another by one pixel or more in the same image were considered as colocalized.

Colocalization analysis for two-color metaphase FISH data for ecDNAs in SNU16 cells described in **Extended Data Figure 8c** was performed using Fiji (version 2.1.0/1.53c)^50^. Images were split into the two FISH colors + DAPI channels, and signal threshold set manually to remove background fluorescence. Overlapping FISH signals were segmented using watershed segmentation. Colocalization was quantified using the ImageJ-Colocalization Threshold program and individual and colocalized FISH signals were counted using particle analysis.

### Immunofluorescence staining and DNA FISH in mitotic cells

For assessing mitotic segregation of ecDNA in GBM39-KT, TR14 and SNU16m1 cells shown in Figure 2, asynchronous cells were grown on poly-L-lysine-coated coverslips (laminin for GBM39-KT). Cells were washed once with PBS and fixed with cold 4% paraformaldehyde (PFA) at room temperature for 10-15 minute. Samples were permeabilized with 0.5% Triton X-100 in PBS for 10 minutes at room temperature and then washed with PBS. Samples were then blocked with 3% BSA in PBS 0.05% Triton X-100 for 30 minutes at room temperature. Samples were incubated in primary antibody (Aurora B Polyclonal Antibody (catalog no. A300-431A; ThermoFisher Scientific), diluted in blocking buffer (1:100–1:200) for either 1 hour at room temperature or overnight at 4 °C. Samples were washed three times in PBS 0.05% Triton X-100. Samples were incubated in secondary antibody, diluted in blocking buffer for 1h at room temperature (all subsequent steps in the dark) and then washed three times in PBS 0.05% Triton X-100. Cells were washed once with PBS and refixed with cold 4% PFA for 20 minutes at room temperature. Cells were washed once with PBS then once with 2× SSC buffer. They were then dehydrated in ascending ethanol concentrations of 70%, 85% and 100% for approximately 2 minutes each. FISH probes (Empire Genomics) were diluted 1:4 in hybridization buffer (Empire Genomics) and added to the sample with the addition of a slide. Samples were denatured at 80 °C for 15-20 minutes and then hybridized at 37 °C overnight in a humid and dark chamber. Samples were then washed with 0.4× SSC then 2× SSC 0.1% Tween 20 (all washes lasting approximately 2 minutes). 4,6-Diamidino-2-phenylindole (DAPI) (100ng/ml) was applied to samples for 10 minutes. Samples were then washed again with 2× SSC 0.1% Tween 20 then 2× SSC. Samples were briefly washed in double-distilled H2O and mounted with ProLong Gold. Slides were sealed with nail polish. Samples were imaged using a DeltaVision Elite Cell Imaging System (Applied Precision) and microscope (model IX-71; Olympus) controlled by the SoftWoRx software v.6.5.2 (Applied Precision) and a 60x objective lens with a CoolSNAP HQ2 camera (Photometrics). Z-stacks were acquired and used to generate maximum intensity projections (ImageJ) for downstream analysis.

For assessing mitotic segregation of oncogene and enhancer ecDNAs in SNU16 cells as shown in Figure 4, cells were seeded onto fibronectin-coated 22×22 coverslips contained in a 6-well culture plate at about 70% confluence. 24 hours after cell seeding, the cells were fixed with 4% PFA and permeabilized with 1× PBS containing 0.25% Triton-X 100. Samples were blocked with 3% BSA-1× PBS for 1 hour at room temperature, followed by primary antibody incubation (Aurora B Kinase antibody; catalog no. A300-431A; Thermo Fisher Scientific) (1:200 in 3% BSA) overnight at 4°C. The sample was washed thrice in 1× PBS followed by incubation with diluted an anti-rabbit Alexa Fluor 647 antibody (1:500 in 3% BSA) for 1 hour at room temperature. The sample is then washed thrice in 1× PBS and fixed with 4% PFA for 20 minutes at room temperature. DNA FISH was performed as described under ‘Metaphase DNA FISH’, with the conditions to heat denaturation changed to 80°C for 20 minutes. Images were acquired on a Leica DMi8 widefield microscope using a 63× oil objective, and each z plane was post-processed by Small Volume Computational Clearing on LAS X prior to generating max projection images.

### Mitotic cell imaging analysis

To quantify fractions of ecDNAs segregated to each daughter cell in pairs of dividing cells as shown in **Figure 2**, ecDNA pixel intensity were quantified from maximum intensity projections using the ImageJ software. ecDNA pixel intensity was measured using the “Integrated Density” measurement from ImageJ. Prior to quantification, background signal from FISH probes was removed uniformly for the entire image until all background signal from the daughter cell nuclei was removed.

To measure fractions of oncogene and enhancer ecDNAs segregated to daughter cells in dividing cells as shown in **Figure 4**, Images were split into the different FISH colors + DAPI channels, and signal threshold set manually to remove background fluorescence using Fiji (version 2.1.0/1.53c)^50^. Overlapping FISH signals were segmented using watershed segmentation. All FISH color channels except DAPI were stacked and ROIs were drawn manually to identify the two daughter cells, after which the color channels were split again and image pixel areas occupied by FISH signals were analyzed using particle analysis. Fractions of ecDNAs in each daughter cell were estimated by fractions of FISH pixels in the given daughter cell.

### Simulations of ecDNA segregation in pairs of daughter cells

To understand how co-segregation dynamics of ecDNAs in dividing cells may affect copy number correlations in daughter cells, we simulated distributions of ecDNA copies among two daughter cells by random sampling using the sample function in R, for which the sample size is the total copy number of an ecDNA species multiplied by two (as a result of DNA replication). For a given fraction of one ecDNA species that co-segregates with the same fraction of another ecDNA species, The corresponding ecDNA copies were randomly distributed among two daughter cells but at the same ratio for both ecDNA species.

To compare observed ecDNA segregation with these simulations given a nonzero frequency of covalent fusions between two ecDNAs such as those between the enhancer and oncogene sequences shown in **Figure 4**, the fraction of fused ecDNAs was treated as co-segregating ecDNAs in the simulations. Thus, for each mitotic immunofluorescence and FISH image collected, the fractions of enhancer ecDNAs, oncogene ecDNAs, and fused enhancer-oncogene ecDNAs were used to simulate 20 segregation events in which a fraction of ecDNAs corresponding to the fused molecules were perfectly co-segregated. The resulting copy number correlations in simulated daughter cells represent the null distribution of ecDNAs explained by covalent fusion alone with no additional co-segregation between distinct ecDNA molecules.

### ATAC-seq

ATAC-seq data for SNU16 were previously published under GEO accession GSE159986^9^. Adapter-trimmed reads were aligned to the hg19 genome using Bowtie2 (2.1.0). Aligned reads were filtered for quality using samtools (version 1.9)^51^, duplicate fragments were removed using Picard’s MarkDuplicates (version 2.25.3), and peaks were called using MACS2 (version 2.2.7.1)^52^ with a q-value cut-off of 0.01 and with a no-shift model.

### ChIP-seq

ChIP-seq data for SNU16 were previously published under GEO accession GSE159986^9^. Paired-end reads were aligned to the hg19 genome using Bowtie2^53^ (version 2.3.4.1) with the --very-sensitive option following adapter trimming with Trimmomatic^44^ (version 0.39). Reads with MAPQ values less than 10 were filtered using samtools (version 1.9) and PCR duplicates removed using Picard’s MarkDuplicates (version 2.20.3-SNAPSHOT). ChIP-seq signal was converted to bigwig format for visualization using deepTools bamCoverage^54^ (version 3.3.1) with the following parameters: --bs 5 --smoothLength 105 --normalizeUsing CPM --scaleFactor 10.

### Evolutionary modeling of ecDNA copy-number framework

EcDNA copy number was simulated over growing cell populations using a forward-time simulation implemented in Cassiopeia^55^ (https://github.com/yoseflab/cassiopeia). All simulations performed in this study were of 2 distinct ecDNA species in a growing cell population. Simulations were parameterized with (i) initial ecDNA copy numbers (initial copy number for ecDNA species *j* is denoted as 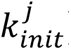); (ii) selection coefficients for cells carrying no ecDNA (*S*_−,−_), both ecDNAs (*S*_+,+_), or either ecDNA (*S*_−,+,’_ or *S*_+,−_; in this study, selection coefficients are treated as constant functions of the types of ecDNA species present in a cell); (iii) a base birth rate (*λ_base_* = 0.5); (iv) and a co-segregation coefficient (*γ*). Optionally, a death rate can also be specified (*μ*).

Starting with the parent cell, a birth rate is defined based on the selection coefficient acting on the cell, *S* ∈ {*S*_−,−_, *S*_−,+’_, *S*_−,+_, *S*_+,+_} as *λ*_1_ = *λ_base_* ∗ (1 + *S*). Then, a waiting time to a cell division event is drawn from an exponential distribution: *t*_b ∼_ exp (−*λ*_1_). When a death rate is also specified, a time to a death event is also drawn from an exponential distribution: *t*_b ∼_ exp (−*μ*). If *t*_b_ < *t*_d_, a cell division event is simulated and a new edge is added to the growing phylogeny with edge length *t*_b_; otherwise, the cell dies and the lineage is stopped. This process will continue until a user-defined stopping condition is specified – either a target cell number (e.g., 1 million) or a target time limit.

During a cell division, ecDNAs are split amongst daughter cells (*d*_1_ and *d*_2._) according to the co-segregation coefficient, *γ*, and the ecDNA copy numbers of the parent cell *p*. In this study, this co-segregation is simulated using two different strategies to determine the effects of co-segregation (see section below entitled “Alternative model of ecDNA co-evolution”). In the following description, let 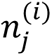 indicate the copy number of ecDNA species *j* in daughter cell *i* and let *N*_j_ indicate the copy number of ecDNA species *j* in the parent cell.

ecDNA species 1 is randomly split distributed to each daughter cell:

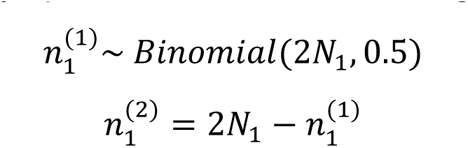

Where *Binomial* is the binomial probability distribution. To simulate co-segregation, for the second ecDNA species, copies are distributed to the daughter cells in proportion to the segregation coefficient *γ* and the copy number of first ecDNA species in each daughter cell:

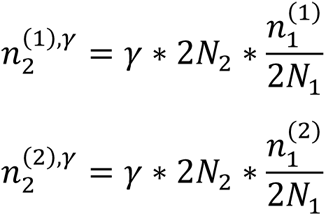

Then, the remainder of copies left over that were not passed with co-segregation are randomly distributed between daughter cells:

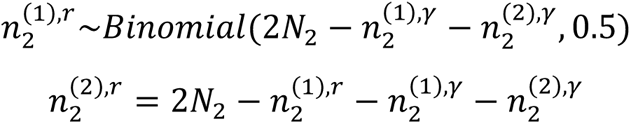

After this simulation, the output is a phylogeny *T* over *l* leaves (denoted by *L*) with ecDNA copy numbers 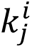 for ecDNA species *j* in leaf *i*.

### Evolutionary modeling of ecDNA co-assortment trends

To simulate the trends of ecDNA copy-number dynamics, we employed the evolutionary modeling framework described previously (see section entitled “Evolutionary modeling of ecDNA copy-number framework”). We used the following fixed parameters: selection acting on individual ecDNA (*S*_−,+_, *S*_+,−_) of 0.2, selection acting on cells without ecDNA (*S*_−,−_) of 0.0, a base birth rate (*λ_base_*) of 0.5, and initial ecDNA copy numbers for both species 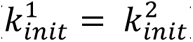 of 5 in the parental cell. We varied co-selection (*S*_+,+’_) and co-segregation (*γ*) between 0 and 1.0 and reported the fraction of cells reporting a copy-number of both ecDNAs above a threshold *m* (by default 1) and the Pearson correlation between ecDNA copy numbers in cells:

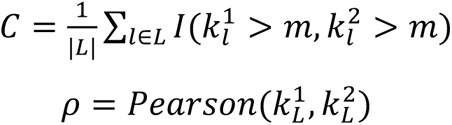

Where 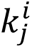 is the copy number of ecDNA species *i* in leaf *l* and 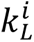 is the vector of copy numbers of ecDNA species *i* across all cells.

For results presented in **Figure 3b-e** and **Extended Figure 4b-d**, we simulated populations of 1 million cells and reported the average co-occurrence and correlation across 10 replicates.

### Inference of evolutionary parameters

Approximate Bayesian Computation (ABC) was used to determine evolutionary parameters in cell line data, specifically selection acting on individual ecDNAs (assumed to be equal between ecDNAs (*S*_−,+_,*S*_+,−_), the level of co-selection (*S*_+,+_), and the co-segregation coefficient (*γ*). Briefly, ABC takes a parameter set *θ* from a prior or proposal distribution and simulates a dataset *y*_0_ from this parameter set. If the simulated dataset matches the observed dataset within specified error tolerance *ε*, then we accept the parameter set and update our posterior distribution *π*(*θ*|*y*_0_). In our case, we defined the priors over each parameter as follows:

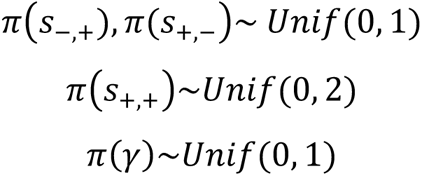

We used the evolutionary model presented above (see section titled “Evolutionary modeling of ecDNA copy-number framework”) to simulate datasets *y*_0_ from the proposed parameter set *θ*, no death rate, a base birth rate *λ_base_*= 0.5, and selection acting on cells without ecDNA *S*_−,−_ = 0.

Here, our goal is to infer a posterior distribution over each evolutionary parameter given single-cell copy-numbers observed from scATAC-seq data in a target cell line, denoted as *y*_obs_ (see above section titled “Paired single-cell ATAC-seq and RNA-seq analysis”). To accomplish this, we chose to derive summary statistics describing the co-occurrence (proportion of cells carrying more than 2 copies of each gene amplified as ecDNA) and the Pearson correlation between the log copy numbers of ecDNAs for guiding our inference, denoted by *C*_obs_ and *ρ*_obs_ respectively. In each round of ABC, we simulated a dataset *y*_0_ of 500,000 cells and compared the summary statistics of this simulated dataset to the observed summary statistics using the following distance function:

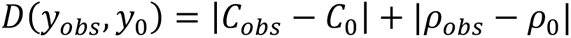

where *C*_0_ and *ρ*_0_ are the simulated co-occurrence and Pearson correlation, respectively. We used an tolerance of *ε* = 0.05 as our target error. Each simulation was initialized with a parental cell with equal copy-number of initial ecDNA (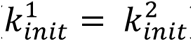): in **Figure 3g** this initial copy number was 5 though alternative initial conditions are explored in **Extended Data Figure 7**. We used the following summary statistics for each cell line: SNU16m1: (*C*_obs_ = 0.99, *ρ*_obs_ = 0.46); TR14: (*C*_obs_ = 0.96, *ρ*_obs_ = 0.26); GBM39-KT: (*C*_obs_ = 0.67, *ρ*_obs_ = 0.36).

The specific implementation of this procedure was performed using a Sequential Monte Carlo scheme (ABC-SMC) using the python package *pyabc* (version 0.12.8). Briefly, this approach performs sequential rounds of inference while computing a weight for the accepted parameters for each iteration. For a more detailed treatment of this procedure, we refer the reader to Sission et al.^56^, Beaumont et al.^57^, Toni et al.^58^ and Lintusaari et al.^59^

### Cell-level co-segregation model of ecDNA co-evolution (Extended Data Figure 5)

Previously, we introduced the co-segregation on the ecDNA element level inside of each cells, where an ecDNA element carrying one species is linked to another element with a probability defined as the co-segregation parameter. Here, we introduce an alternative model, where ecDNA co-segregation is implemented on a cellular level. In each cell division, if a cell is chosen for proliferation, the number of ecDNA copies in that cell are doubled. We first have the randomly segregation of both ecDNA species following a binomial distribution seperately, and then pair those with high copy numbers into the same daughter cells with a probability *γ* ∈ [0,1]. More precisely, *γ* describes the likelihood of extreme copy number correlation, and 1 − *γ* describes the likelihood of extreme copy number anticorrelation. If *γ* = 0.5, it is related to unbiased likelihood for both extreme scenarios, and it results in the modeling of standard random ecDNA proliferation without co-segregation.

In this model, the population growth is also modeled as a birth-death stochastic process and implemented by a standard Gillespie algorithm^4^. We starting from a small initial population (a single cell or three cells) carrying a certain amount of ecDNA elements and recording the exact number of ecDNA copies for each cell through the simulation. Cells are chosen randomly but proportional to their fitness (1+*S*) for proliferation, where *S* is the selection coefficient. Neutral proliferation is defined compared to fitness of cells without ecDNA (*S* = 0). If there is a fitness effect by carrying ecDNA, *S* >0. For simplicity, in our models, we give a fixed selection coefficient for cells carrying either ecDNA and vary the selection coefficient for cells with both ecDNA to investigate the impact of co-selection in ecDNA co-evolution. For reporting, we discretise the population into three subpopulations, named pure, mix and free (no) ecDNA cells (**Figure 3h**), which represent cells carrying just one type of ecDNA, both types or no ecDNA at all respectively. For the results presented in **Extended Data Figure 5**, we simulated populations of 10,000 cells and reported summary statistics across 500 replicates.

### Evolutionary modeling of drug intervention

The evolutionary model described previously (see section titled “Evolutionary modeling of ecDNA copy-number framework”) was used to evaluate the effect of Pemigatinib treatment on SNU16m1 cells. To do so, we modified the framework to allow for a “burn-in” period to simulate population growth without drug and then introduced a perturbation to selection coefficients at a defined time point.

Specifically, we allowed the cell population to grow to 5000 cells under the following conditions: base birth rate (*λ_base_*) of 0.5, a death rate (*μ*) of 3, an initial ecDNA copy number for both species (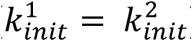) of 10, the following selection coefficients: *S*_−,−_ = 0; *S*_−,+_ = 0.15; *S*_+,−_ = 0.15; *S*_+,+_ = 0.8 (here, let cells carrying only *FGFR2* ecDNA be denoted by *S*_+,−_ and cells only carrying *MYC* ecDNA by *S*_−,+_). We allowed co-segregation to vary 0 and 1.

After a population of 5000 cells was obtained, we simulated Pemigatnib treatment by changing the selection pressures acting on cells. Specifically, we set *S*_+,+_ = *S*_+,−_ ∈ {0, −0.1, −0.2, −0.3, −0.4, −0.5}. We then simulated 500,000 cells from the pre-treatment group of 5,000 cells while maintaining the same values for *γ*, *μ*, *λ_birth_*, and *S*_−,−_. For time-dependent functions of copy-number reported in **Figure 4i**, mean copy-number of both ecDNA species were computed in time bins of 5 up until the introduction of Pemigatinib and bins of 1 afterwards.

### Evolutionary modeling of enhancer-only ecDNA

To explore the evolutionary principles of enhancer-only ecDNA, we used the previously described evolutionary model (see above “Evolutionary modeling of ecDNA copy-number framework”) without death and fixed the following evolutionary parameters: *S*_+,−_ = 0.2, *S*_−,+_ = 0, *λ_base_* = 0.5, and 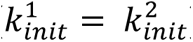 = 5. We simulated 10 replicates of 1M cell populations a modulated co-selection coefficient *S*_+,+_ from [0, 1] and co-segregation coefficient *γ* from [0,1]. In Figure 4, we report the distribution of co-occurrence summary statistics *C* across these 10 replicates.

### Nanopore sequencing of SNU16 genomic DNA

Genomic DNA from approximately 2 million SNU16 cells was extracted using the MagAttract HMW DNA Kit (Qiagen 67563) and prepared for long-read sequencing using a Ligation Sequencing Kit V14 (Oxford Nanopore Technologies SQK-LSK114) according to the manufacturer’s instructions. Libraries were sequenced on a PromethION (Oxford Nanopore Technologies) using a 10.4.1 flow cell (Oxford Nanopore Technologies FLO-PRO114M).

Basecalling from raw POD5 data was performed using Dorado (Oxford Nanopore Technologies, version 0.2.1+c70423e). Reads were aligned using Winnowmap2^60^ (version 2.03) with the following parameters: -ax map-ont. Structural variants were called using Sniffles^61^ (version 2.0.7) using the following additional parameters: --output-rnames.

### Pemigatinib treatment of SNU16m1 and COLO320-DM cell lines

SNU16m1 cells were treated with 1 *μ*M or 5 *μ*M Pemigatinib (Selleckchem: INCB054828), or with an equal volume of PBS. COLO320-DM cells were treated with 5 uM Pemigatinib or an equal volume of PBS. Fresh Pemigatinib was replenished approximately every 7 days. Approximately 2 million SNU16m1 cells were sampled from each condition at day 0, 6 and 20, genomic DNA was extracted using the Quick DNA MiniPrep kit (Zymo Research; D0325), and subjected to WGS (see above, section entitled “WGS”). Approximately 2 million COLO320-DM cells were sampled at day 20 and genomic DNA was prepared for sequencing using the same procedure as above.

### Nutlin-3a treatment of TR14 cells and interphase DNA FISH

175,000 TR-14 cells were seeded per well in 12-well plates. Cells were treated either with 0.1% DMSO or with 1 µl Nutlin-3a (Sigma Aldrich, SML0580) for 6 days, without an additional wash-out period.

Samples were fixed using Carnoy’s Solution (3:1 Methanol:Acetic Acid). Fixed samples on coverslips or slides were briefly equilibrated in 2× SSC buffer. They were then dehydrated in ascending ethanol concentrations of 70%, 90% and 100% for approximately 2 minutes each. FISH probes were diluted in Probe Hybridization Buffer and added to the sample with the addition of a coverslip or slide. Samples were denatured at 78 °C for 5 min and then hybridized at 37 °C overnight in a humid and dark chamber. Samples were washed twice in 0.4X SSC with 0.3% IGEPAL CA-630 for 2 min with agitation for the first 10-15 seconds. They were then washed once in 2X SSC with 0.1% IGEPAL CA-630 at room temperature for 2 minutes, again with agitation for the first 10-15 sec. 4,6-Diamidino-2-phenylindole (DAPI) (100 ng ml−1) was applied to samples for 10 minutes. Samples were then washed again with 2× SSC and mounted with ProLong Antifade Mountant.

FISH and microscopy was carried out in the same manner as TR14 was processed as described above (see section entitled “Metaphase DNA FISH image analysis”). Normalized copy numbers *n*_*i*_ for ecDNA *i* were computed from raw copy numbers *c*_*i*_ as:

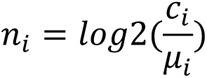

where *μ*_*i*_ is the mean copy number of ecDNA *i* in the DMSO control condition. Statistical significance was assessed with the Wilcoxon rank sums test.

## Data Availability

WGS data from bulk SNU16 cells were previously generated (SRR530826, Genome Research Foundation). Paired single-cell ATAC-seq and RNA-seq data for COLO320DM were generated previously and published under GEO accession GSE159986. TR14 scCircle-seq data were deposited in the European Genome-phenome Archive (EGA) under the accession number: EGAS00001007026. CRISPR-CATCH sequencing data integrated from previous studies were deposited in NCBI Sequence Read Archive (SRA) under BioProject accessions PRJNA670737 and PRJNA777710. ATAC-seq and ChIP-seq data for SNU16 were previously published under GEO accession GSE159986.

## Code Availability

The ecDNA evolutionary modelling framework used in this study is publicly available through Cassiopeia^40^ at https://github.com/YosefLab/Cassiopeia. AmpliconClassifer is available at https://github.com/jluebeck/AmpliconClassifier.

## Materials & Correspondence

Correspondence and requests for materials should be addressed to Howard Y. Chang and Paul S. Mischel.

