## Supplementary Information for "Coordinated inheritance of extrachromosomal DNA species in human cancer cells"

**Table of Contents**

**Supplementary Table 1:** CRISPR guide RNA sequences.

### Supplementary Tables

#### Supplementary Table 1.

| Guide RNA sequence | Guide RNA information |
|---|---|
| CCAGCAATCGTTAACCACTG | *MYC* guide 3 |
| CTTCGGGGAGACAACGACGG | *MYC* guide 5 |
| GTGATATTTGAACCGCCCTG | *MYC* guide 7 |
| GTCTTGCTAGACTATACCAG | *MYC* guide 19 |
